# Hierarchical assembly and functional resilience of the mammalian RNA exosome

**DOI:** 10.1101/2025.03.14.643291

**Authors:** Tsimafei Navalayeu, Nikolaus Beer, Monika Bebjaková, Robert W. Kalis, Karel Stejskal, Gabriela Krššáková, Nina Fasching, Veronika A. Herzog, Niko Popitsch, Elisabeth Roitinger, Johannes Zuber, Stefan L. Ameres

**Affiliations:** Max Perutz Labs, Vienna BioCenter (VBC), Dr.-Bohr-Gasse 9, A-1030, Vienna, Austria; University of Vienna, Max Perutz Labs, Department of Biochemistry and Cell Biology, Dr.-Bohr-Gasse 9, A-1030 Vienna, Austria; Vienna BioCenter PhD Program, a Doctoral School of the University of Vienna and the Medical University of Vienna, A-1030 Vienna, Austria; Institute of Molecular Biotechnology of the Austrian Academy of Sciences (IMBA), Vienna BioCenter (VBC), A-1030 Vienna, Austria; Research Institute of Molecular Pathology (IMP), Vienna BioCenter (VBC), 1030 Vienna, Austria; Medical University of Vienna, Vienna BioCenter (VBC), Vienna, Austria

## Abstract

Most eukaryotic proteins assemble into multisubunit complexes that coordinate essential cellular functions, yet the principles governing their assembly and proteostatic control remain largely undefined. Here, we systematically dissect the cellular assembly and functional organization of the RNA exosome, an essential ribonucleolytic complex, using an inducible dual-guide CRISPR/Cas9 system in mouse embryonic stem cells. We reveal a sequential assembly pathway where Exosc2, Exosc4, and Exosc7 initiate complex formation, facilitating the incorporation of barrel and cap subunits in a defined hierarchy. Unlike other structural subunits, the terminally incorporated cap subunit Exosc1 is dispensable for cell viability, revealing a modular, functionally resilient architecture. We demonstrate that orphan subunits are selectively degraded via the ubiquitin-proteasome system, enforcing stringent quality control over RNA exosome biogenesis. These findings establish a framework for decoding the assembly logic of essential macromolecular machines and uncover previously unrecognized plasticity in the composition and function of the RNA exosome.

## Introduction

The biogenesis and stability of multiprotein complexes are fundamental to cellular function. Most eukaryotic proteins act within structured assemblies that coordinate essential processes such as transcription, translation and RNA metabolism^1–3^. The accurate assembly of these complexes is tightly regulated, with dedicated quality control mechanisms recognizing and eliminating orphan subunits that fail to integrate^4,5^. Despite the prevalence of multiprotein complexes, their assembly pathways and proteostatic dependencies remain poorly understood. While detailed studies have elucidated the stepwise biogenesis of large protein assemblies such as the ribosome and proteasome^5–9^, how RNA-processing complexes form and how their stability is maintained in cells remains largely obscure.

The RNA exosome is a conserved ribonucleolytic complex that regulates the processing and degradation of most coding and non-coding RNA species across all eukaryotic domains^10^. In mammals, the RNA exosome consists of a catalytically inert core, comprised of a cap formed by Exosc1, Exosc2, and Exosc3, and a barrel-module composed of Exosc4, Exosc5, Exosc6, Exosc7, Exosc8, and Exosc9^11^. This RNA exosome core provides a scaffold for 3’-5’ exoribonucleases, namely Exosc10 and Dis3 in the nucleus, and Dis3L in the cytoplasm^12–14^. The RNA exosome is crucial for clearing products of pervasive transcription, such as Promoter Upstream Transcripts (PROMPTs) and enhancer RNAs (eRNAs) in the nucleus, as well as for quality control and degradation of mRNA in the cytoplasm^15–20^. In addition to its RNA decay function, the RNA exosome contributes to the processing and maturation of various non-coding RNA species, including small nuclear RNA (snRNA), small nucleolar RNA (snoRNA) and ribosomal RNA (rRNA)^18,21–23^. Given its central role in RNA metabolism, it is essential to define how the RNA exosome assembles in a cellular context and how its integrity is monitored.

Several *in vitro* reconstitution approaches of the yeast and mammalian RNA exosome provided important insights into its architecture and inter-subunit interactions and indicated that pre-formed dimeric or trimeric subcomplexes may coalesce to form the RNA exosome^11,12,24–26^. However, whether exosome assembly follows a defined sequence *in vivo* or whether subunits can integrate independently remains unclear. It is also unknown whether the incorporation of all subunits is an obligatory requirement for RNA exosome function or whether partially assembled complexes can retain activity.

Genetic perturbation studies suggest that the individual contributions of RNA exosome subunits to complex function may differ. While most core components are required for cell viability in diverse model systems ^15–20^, Exosc1 appears to be an exception. In *Trypansoma brucei*, depletion of all RNA exosome subunits except Exosc1 resulted in severe growth defects^27^. In mice, comparison of Exosc1 and Exosc2 null embryos revealed substantial phenotypic differences. Exosc2 mutants exhibit impaired uterine implantation at embryonic day 3.5 to 4 and cannot be recovered at post-implantation stages. In contrast, Exosc1 null embryos are able to progress further and form an egg-cylinder, but fail to initiate gastrulation and develop morphologically past embryonic day 5.5^28^. These findings raise the possibility that Exosc1 is not strictly required for RNA exosome function and that a structurally incomplete complex may still retain activity. However, the physiological and molecular consequences of Exosc1 loss remain unresolved, and its potential role in exosome assembly and stability has not been systematically examined.

Current evidence suggests that many multiprotein complexes are assembled in a manner tightly linked to proteostatic surveillance, where orphan or misassembled subunits are selectively degraded to maintain complex integrity^5–9^. Whether similar principles apply to RNA exosome biogenesis, however, remains unknown. While few proteostatic interdependencies were reported for the RNA exosome subunits^13,16,19,29–34^, it is unclear whether their turnover is a direct consequence of failed assembly or if additional regulatory mechanisms actively govern exosome stability. Moreover, it is unknown whether RNA exosome formation requires stepwise stabilization of specific intermediates or if it follows a more flexible incorporation pathway that allows dynamic subunit exchange. Resolving these questions is critical for understanding how the RNA exosome is maintained as a functional entity under physiological and stress conditions.

Here, we systematically investigate RNA exosome assembly and proteostatic regulation in mouse embryonic stem cells (mESCs). Using inducible dual-guide CRISPR/Cas9 genome engineering alongside acute auxin-inducible degron (AID) protein depletion, we define a sequential assembly hierarchy and uncover mechanisms that regulate RNA exosome stability. We show that orphan subunits are selectively degraded via the ubiquitin-proteasome system, linking RNA exosome biogenesis to proteostatic surveillance. We further demonstrate that Exosc1, an integral member of the RNA exosome cap structure, is dispensable for RNA exosome function, revealing an unexpected resilience in exosome architecture. These findings provide a framework for understanding the cellular principles of RNA exosome assembly and expand our knowledge of how multiprotein complexes are dynamically formed, stabilized, and regulated in cells.

## Results

### A dual guide iCas9 system probes the essential cellular function of RNA exosome subunits in mESCs

To systematically assess the role of individual Exosc proteins under conditions that take into account the essential role of the RNA exosome, we established a doxycycline (Dox)-inducible Cas9 system (iCas9) in mESCs (Figure 1A and S1A)^35^. For gRNA design we utilized the VBC score, which considers gRNA activity, frequency of indel formation as well as amino acid composition and conservation of the targeted region to increase knockout efficiency and penetrance across a cell population^36^. To enhance protein depletion efficiency, we employed a dual gRNA targeting strategy that mitigates the potential poor editing efficiency of single gRNAs and diminishes reversions emerging from in-frame repair of single sites (Figure 1A). We functionally validated this iCas9 system by targeting a surface marker (CD9) and yellow fluorescent protein (YFP), confirming consistent and complete protein depletion only in the presence, but not in the absence of Dox (Figures S1B). Upon introducing dual gRNAs targeting each one of the nine structural core components (Exosc1 to Exosc9; Figure 1B) of the RNA exosome or its associated catalytic subunits (Exosc10, Dis3, and Dis3L; Figure 1B), followed by Cas9 induction, we observed a progressive and consistent reduction in target protein abundance by Western blot analyses, reaching maximal depletion (i.e. ≤ 10%) after 72 hours (Figure 1C and D). This was consistent with previously reported RNA exosome protein half-live measurements in murine cells^37^.

**Figure 1.**
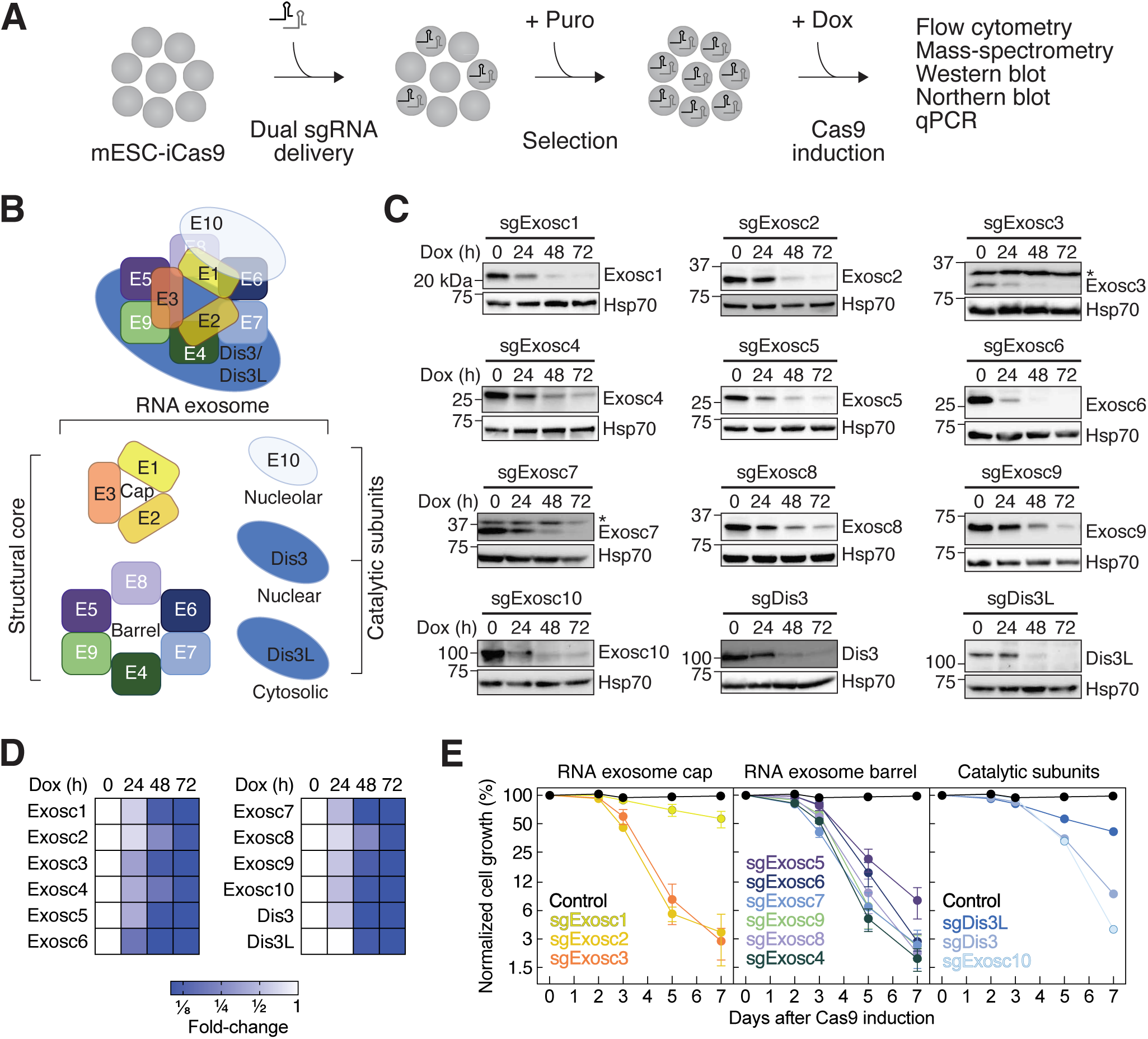
A dual-guide iCas9 system probes the essential cellular function of RNA exosome subunits in mESCs. **(A)** Outline of the experimental approach. Dual sgRNAs targeting RNA exosome subunits were delivered to iCas9 mESCs, followed by Puromycin (Puro) selection and Cas9 induction with doxycycline (Dox). **(B)** Schematic of the mammalian RNA exosome, highlighting core (cap and barrel) and catalytic subunits.”E” denotes Exosc subunits (e.g. E1 = Exosc1). **(C)** Western blot analysis of iCas9 mESCs expressing dual sgRNAs targeting the indicated RNA exosome component and treated with Dox for 0h, 24h, 48h and 72h. Hsp70 serves as a loading control. Asterisk (*) indicates a non-specific band. **(D)** Western blot quantifications from (C) are shown as a heatmap. Protein levels were normalized to Hsp70 and expressed relative to the 0h time point (n = 2 biological replicates). **(E)** Competitive proliferation assays of inducible Exosc knockout and control Olfr10 (control) mESCs relative to a parental control cell line. Data were normalized to Day 0 and presented as mean ± SD (n = 3 biological replicates). See also **Figure S1**.

To explore the physiological consequences following the loss of individual RNA exosome components we conducted a competitive proliferation assay by mixing parental iCas9-mESCs with dual gRNA (and mCherry) expressing cells in the presence of Dox (Figure 1E). To exclude potential target-independent adverse effects of Cas9 and/or dual gRNA expression on cell growth, we utilized as a control a gRNA pair targeting the olfactory receptor 10 (Olfr10) gene, which is not normally expressed in mESCs. In contrast to Olfr10-targeting, which caused no discernible proliferation defects, significant growth impairment was observed upon the depletion of most structural RNA exosome core subunits, except for Exosc1. Upon Exosc1 depletion, the fraction of knockout cells merely dropped to ∼50% after 7 days of Cas9 induction, in contrast to the less than 10% of gRNA-expressing cells remaining after the depletion of each other RNA exosome core component. Targeting Exosc1 with additional pairs of gRNAs exhibited similarly weak proliferation defects, and CRISPR-based viability screens in human cancer cell lines indicate that Exosc1 knockout has a much lower impact on cell survival than other core subunits^38–40^, confirming its subordinate role in sustaining cellular proliferation (Figures S1C and D).

To benchmark the proliferation defects observed upon depletion of individual structural RNA exosome core subunits by iCas9 against a complementary acute protein depletion strategy, we introduced an auxin-inducible degron (AID) tag at the endogenous N-terminus of Exosc3 or the C-terminus of Exosc6 in Tir1-expressing mESCs^41^. Tagging Exosc3 and Exosc6 with AID prompted only minor changes in their protein abundance in the absence of IAA and did not affect proliferation, confirming that genetic engineering of Exosc6 and Exosc3 in mESCs did not perturb RNA exosome integrity (Figure S1E-F). In contrast, administration of indole-3-acetic acid (IAA) resulted in a rapid depletion of AID-Exosc3 and Exosc6-AID proteins (Figure S1E) and induced substantial growth defects (Figure S1F), consistent with the inducible knockout (iKO) experiments described above (Figure 1E).

Lastly, we performed growth competition experiments following the inducible knockout of the RNA exosome-associated catalytic subunits (Figure 1E). Depletion of the nuclear catalytic subunits Exosc10 and Dis3 resulted in pronounced growth impairment, while the loss of the cytoplasmic catalytic subunit Dis3L led to moderate proliferation defects, in agreement with previous reports^18,42,43^.

In summary, we established and benchmarked a dual guide iCas9 system in mESCs to show that the depletion of individual RNA exosome subunits substantially impairs cell proliferation, while Exosc1 and Dis3L play a subordinate role.

### Selective decay of orphan exosome subunits by the ubiquitin-proteasome system reveals an ordered cellular assembly of the RNA exosome core complex

During cellular assembly, protein complexes are frequently surveilled to detect and remove unassembled orphan subunits^4,5^. We systematically investigated whether the loss of individual RNA exosome subunits affects the overall integrity of the RNA exosome by monitoring reciprocal protein depletion. To this end, we targeted each structural component of the RNA exosome (i.e. Exosc1 to Exosc9) for 24h, 48h, and 72h by iCas9, followed Western blot analyses of every RNA exosome subunits (Figure 2A, S1G-O). To quantitatively characterize these co-destabilization patterns, we analysed samples subjected to 72h of inducible knockout using mass-spectrometry (Figure 2B, S1S and Table S3). Indeed, we observed specific patterns of exosome protein co-destabilization that inform on a stepwise cellular assembly of the RNA exosome.

**Figure 2.**
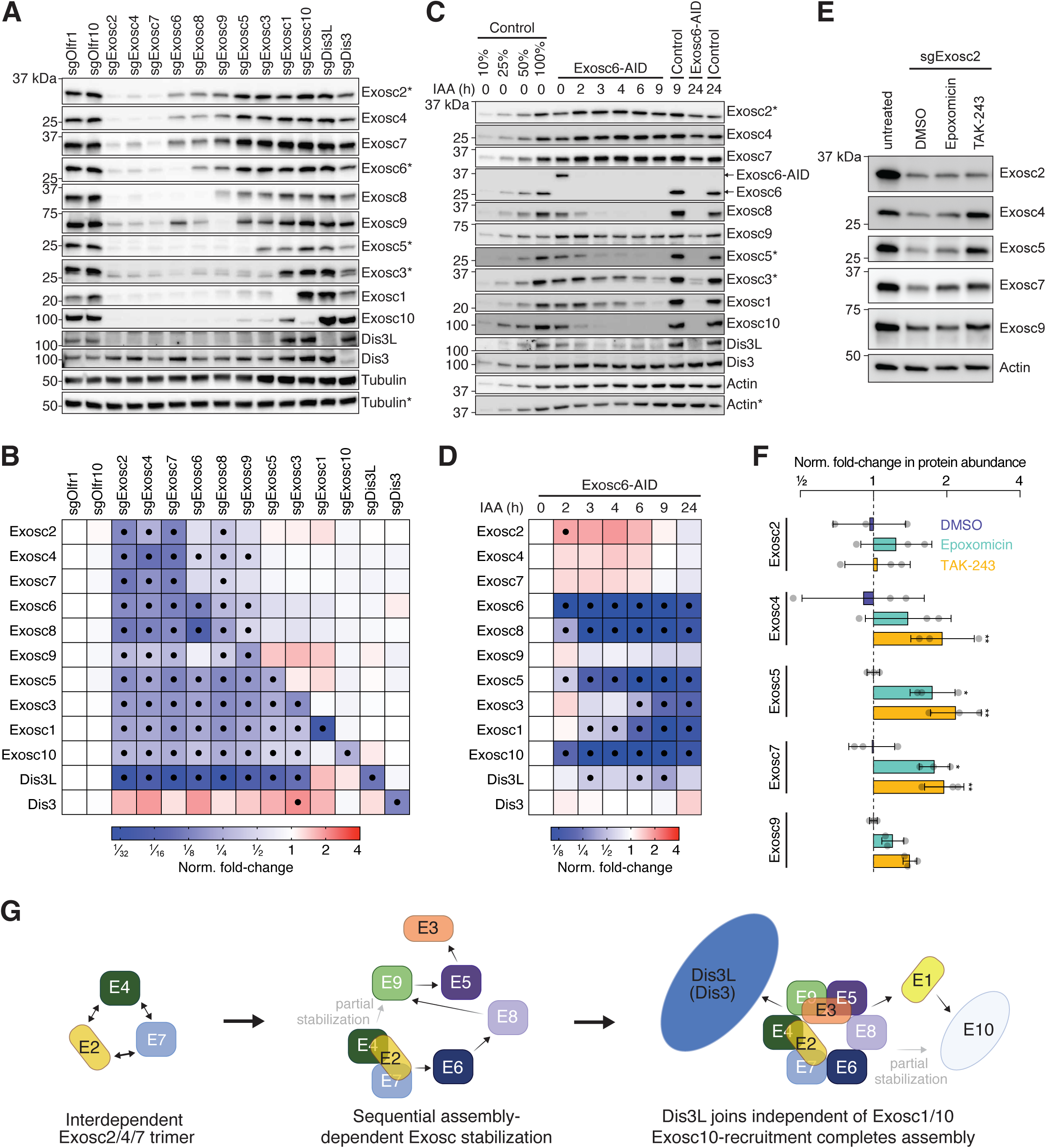
Proteolytic degradation of orphan subunits by the ubiquitin-proteasome system reveals an ordered cellular assembly of the RNA exosome. **(A)** Western blot analysis was performed on iCas9 mESCs harboring dual sgRNAs targeting the indicated RNA exosome components or Olfr1 and Olfr10 (controls), treated with Dox for 72h. Tubulin serves as a loading control. Asterisks (*, **) indicate matching membranes. **(B)** Relative abundance of RNA exosome subunits was assessed 72h after induction of sgRNAs targeting the indicated Exosc, normalized to sgOlfr1 control. Protein abundance was determined by shotgun or targeted mass-spectrometry. Values are shown relative to sgOlfr1 control sample. Statistical significance was determined using two-way ANOVA with Holm-Šídák multiple comparison correction. Black dots indicate instances where log_2_FC < −1 or log_2_FC > 1 and p_adj_ < 0.05. **(C)** Auxin-inducible degradation of Exosc6 was analyzed in Exosc6-AID mESCs treated with IAA for the indicated time in hours (h). Actin serves as a loading control. Asterisk (*) refers to matching membranes. **(D)** Western blot quantifications from (C) were normalized to Actin and expressed relative to untreated Exosc6-AID mESCs (n = 2 biological replicates). Statistical significance was determined using two-way ANOVA with Holm-Šídák multiple comparison correction. Black dots indicate instances where p_adj_ < 0.05. **(E)** Western blot analysis was performed on iCas9 mESCs harboring dual sgRNAs targeting Exosc2 treated with Dox for 42h, followed by 6h of DMSO, Epoxomicin and TAK-243 treatment. Actin serves as a loading control. **(F)** Western blot quantifications from (E) were normalized to Actin and expressed relative to DMSO control (n = 3 biological replicates). Statistical significance was determined using two-way ANOVA with Holm-Šídák multiple comparison correction. (* p_adj_ < 0.05, ** p_adj_ < 0.01, *** p_adj_ < 0.001, **** p_adj_ < 0.0001). **(G)** Model of the cellular assembly of RNA exosome. See text for details. See also **Figure S1** and **Table S3**.

First, depletion of Exosc2, Exosc4 or Exosc7 resulted in strong co-destabilization of all RNA exosome core components (Figure 2A, B and S1G-I). At the same time, these subunits remained stable in knockouts of the other RNA exosome core subunits, suggesting that Exosc2-4-7 form a stable trimeric complex that initiates RNA exosome assembly (Figure S1J-O). Consistently, structural studies show extensive interaction surfaces (∼2000 Å^2^ each) among these three subunits^11^.

Depletion of other RNA exosome core subunit triggered selective proteostatic responses, consistent with a stepwise association of Exosc6, Exosc8, Exosc9, Exosc5, Exosc3, and Exosc1 with the Exosc2/4/7 trimer. Exosc6 depletion strongly impacted the stability of Exosc8, Exosc5, Exosc3, and Exosc1, while Exosc2, Exosc4, Exosc7 and Exosc9 were less impacted (Figure 2A, B and S1J). Given Exosc6’s dependence on Exosc2, Exosc4, and Exosc7 for stability (Figure S1G-I), we concluded that Exosc6 incorporates after Exosc2/4/7 by associating with Exosc7, which forms extensive contacts (2700 Å^2^) with Exosc6 in structural analyses^11^.

Exosc8 depletion affected the stability of Exosc9, Exosc5, Exosc3, Exosc1, while exerting a weaker impact on Exosc2, Exosc4, Exosc7, or Exosc6, suggesting that Exosc8 joins after the formation of the Exosc2/4/7/6 subcomplex (Figure 2A, B and S1K). Consistent with this model, structural data show that Exosc8 interacts with Exosc6 but does not directly contact Exosc2/4/7^11^.

Notably, Exosc9 was only partially destabilized upon depletion of Exosc6 and Exosc8, suggesting a weak protective association with Exosc2/4/7 (Figure 2A, B and S1J and K). However, targeting Exosc9 by iCas9 largely destabilized Exosc5, Exosc3, and Exosc1 (Figure 2A, B and S1L). Structural data indicate that Exosc9 directly contacts Exosc4, suggesting it joins the complex through Exosc4 binding but only after the formation of a Exosc2/4/7/6/8 subcomplex^11^.

Targeting Exosc5 resulted in co-depletion of Exosc3 and Exosc1 (Figure 2A, B and S1M), indicating that Exosc5 integrates after the formation of the Exosc2/4/7/6/8/9 subcomplex, completing the ring formed by the barrel subunits of the RNA exosome. Consistently, Exosc5 shares substantial interaction surfaces with Exosc9 (3800 Å^2^) and Exosc8 (2400 Å^2^)^11^.

Exosc3 depletion resulted in co-depletion of Exosc1 (Figure 2A, B and S1N), suggesting it incorporates after the formation of the Exosc2/4/7/6/8/9/5 subcomplex. This finding is surprising, given that structural data show minimal direct contact between Exosc3 and Exosc1^11^. Thus, Exosc3 may stabilize Exosc1 indirectly by enforcing its interaction with Exosc6 (∼1400 Å^2^) and Exosc8 (∼300 Å^2^), potentially by promoting a cap-binding competent conformation of the barrel through direct contacts with Exosc9 and Exosc5.

Finally, Exosc1 depletion did not affect any other core subunit (Figure 2A, B and S1O), confirming its role as the final component of RNA exosome core assembly, where it directly interacts with Exosc8 and Exosc6^11^. Overall, our observations indicate that depletion of RNA exosome subunits arrests complex assembly at specific stages, while unassembled subunits undergo proteolytic degradation.

To explore the temporal relationship between RNA exosome integrity and proteostatic control, we performed acute protein depletion experiments. We assessed core RNA exosome protein abundance in Exosc6-AID-or AID-Exosc3-engineered Tir1-mESCs at multiple time points (up to 24 h) after IAA treatment using Western blot analyses (Figure 2C, D and S1T, U). Similar to our iCas9 observations, acute depletion of Exosc6 – whose protein levels drop to undetectable levels within 2h of IAA treatment – resulted in significant co-depletion of Exosc8, Exosc5, Exosc3, and Exosc1 as early as 2h post-treatment (Figure 2C and D). Notably, Exosc2, Exosc4, Exosc7 and Exosc9 levels decreased only after the prolonged (24h) depletion of Exosc6 (Figure 2C, D). These results align with the co-destabilization pattern observed upon iCas9 targeting of Exosc6 and support the model that, once the Exosc2/4/7 intermediate is assembled, extensive interactions of Exosc9 with Exosc4 (∼2900 Å^2^) stabilize Exosc9 even in a truncated RNA exosome core complex^11^. Furthermore, depletion of AID-Exosc3 by IAA treatment, which led to the depletion of Exosc3 to undetectable levels within 3 h, affected only Exosc1 levels among all investigated RNA exosome core components, confirming Exosc3 as the penultimate component in RNA exosome assembly (Figure S1T and U). Collectively, these data validate the proposed stepwise assembly model and suggest that selective perturbations in RNA exosome integrity rapidly activate proteostatic mechanisms that eliminate orphan RNA exosome core proteins.

Finally, we investigated the role of the ubiquitin-proteasome system in the turnover of orphan RNA exosome core components. To do so, we depleted Exosc2 by iCas9 for 42 hours, followed by proteasome inhibition with Epoxomicin^44^ or inhibition of the main E1-ubiquitin ligase (Uba1) through TAK-243 for 6 hours^45^. Western blot analyses revealed stabilization of orphan RNA exosome subunits Exosc4, Exosc5, Exosc7 and Exosc9, but not the genetically targeted Exosc2, upon Epoxomycin or TAK-243 treatment compared to a DMSO control (Figure 2E and F).

In summary, our experiments demonstrate that RNA exosome integrity is actively monitored by proteostatic control mechanisms that recognize and degrade orphan core components via the ubiquitin-proteasome system. By integrating these findings with published structural studies^11^, we propose a model for RNA exosome core assembly (Figure 2G): Exosc2, Exosc4, and Exosc7 form an interdependent heterotrimer that serves as the foundation for the sequential incorporation of Exosc6, Exosc8, Exosc9, Exosc5, Exosc3, and finally, Exosc1.

### Catalytic subunits associate with the RNA exosome core at distinct assembly steps

We next examined the proteostatic relationship between structural and catalytic subunits of the RNA exosome complex. When Dis3L, Dis3, or Exosc10 were depleted using iCas9, we did not detect any changes in protein levels of RNA exosome core components (Figure 2A, B, and S1P, Q, and R). Hence, catalytic components do not play a direct role in the assembly of the RNA exosome core.

Depleting any RNA exosome core component, except Exosc1, using iCas9, as well as acutely depleting Exosc6 or Exosc3 using AID, led to substantial co-depletion of the cytoplasmic catalytic subunit Dis3L (Figure 2A-D, and S1G-O, T, and U). These results align with structural data from yeast and human, where Rrp44 (a homolog of murine Dis3L/Dis3) associates with the RNA exosome core via its PIN domain and through Rrp45 (Exosc9)-Rrp41 (Exosc4), with additional contacts from a long N-terminal *β*-hairpin wedged between Rrp41 (Exosc4) and Rrp42 (Exosc7)^46–48^. Hence, Dis3L presumably associates with the cytoplasmic RNA exosome core at a late assembly stage but independently of Exosc1.

In contrast, Dis3, the nuclear counterpart of Dis3L, remained stable under all conditions except when directly depleted using iCas9 (Figure 2C, D and S1G-R), suggesting that Dis3 is not subject to assembly-dependent proteostatic control.

Furthermore, iCas9-mediated depletion of any RNA exosome core component, as well as acute protein depletion of both Exosc6 and Exosc3 by AID, resulted in substantial co-depletion of the nuclear catalytic subunit Exosc10 (Figure 2A-D and S1G-O, T, and U). This implies that Exosc10 stability strongly depends on a fully assembled RNA exosome core. Consistent with this observation, Exosc10/Rrp6 associates with the RNA exosome core via interactions with Exosc1/Csl4, Exosc6/Mtr3, and Exosc8/Rrp43^46,47^.

In summary, our findings show that, unlike Dis3, both Dis3L and Exosc10, are subject to orphan protein decay, which is prevented by their association with either a late-stage or a fully assembled RNA exosome core complexes, respectively.

### RNA exosome assembly intermediates form biochemically tractable subcomplexes

The selective proteolytic degradation of RNA exosome components in perturbation settings suggests a model for the cellular assembly of the RNA exosome that predicts the formation of biochemically tractable subcomplexes (Figure 2G). To investigate this, we generated a cell line with ectopic expression of Exosc2-mCherry-V5 (Figure S2A). Notably, expression of Exosc2-mCherry-V5 led to the destabilization of endogenous Exosc2 (Figure S2B), a phenomenon previously observed in *Trypanosoma brucei* and consistent with the orphan protein decay described in this study^49^. In contrast to the V5-mCherry control, immunoprecipitation of Exosc2-mCherry-V5 recovered all RNA exosome subunits, but not the control Tubulin, validating the effective and specific recovery of fully assembled RNA exosome complexes (Figure 3A). Note, that Dis3 was hardly detectable upon Exosc2-V5 immunoprecipitation, consistent with previously published experiments and perhaps pertaining to the absence of its assembly-dependent proteostatic control^13,14,50^.

**Figure 3.**
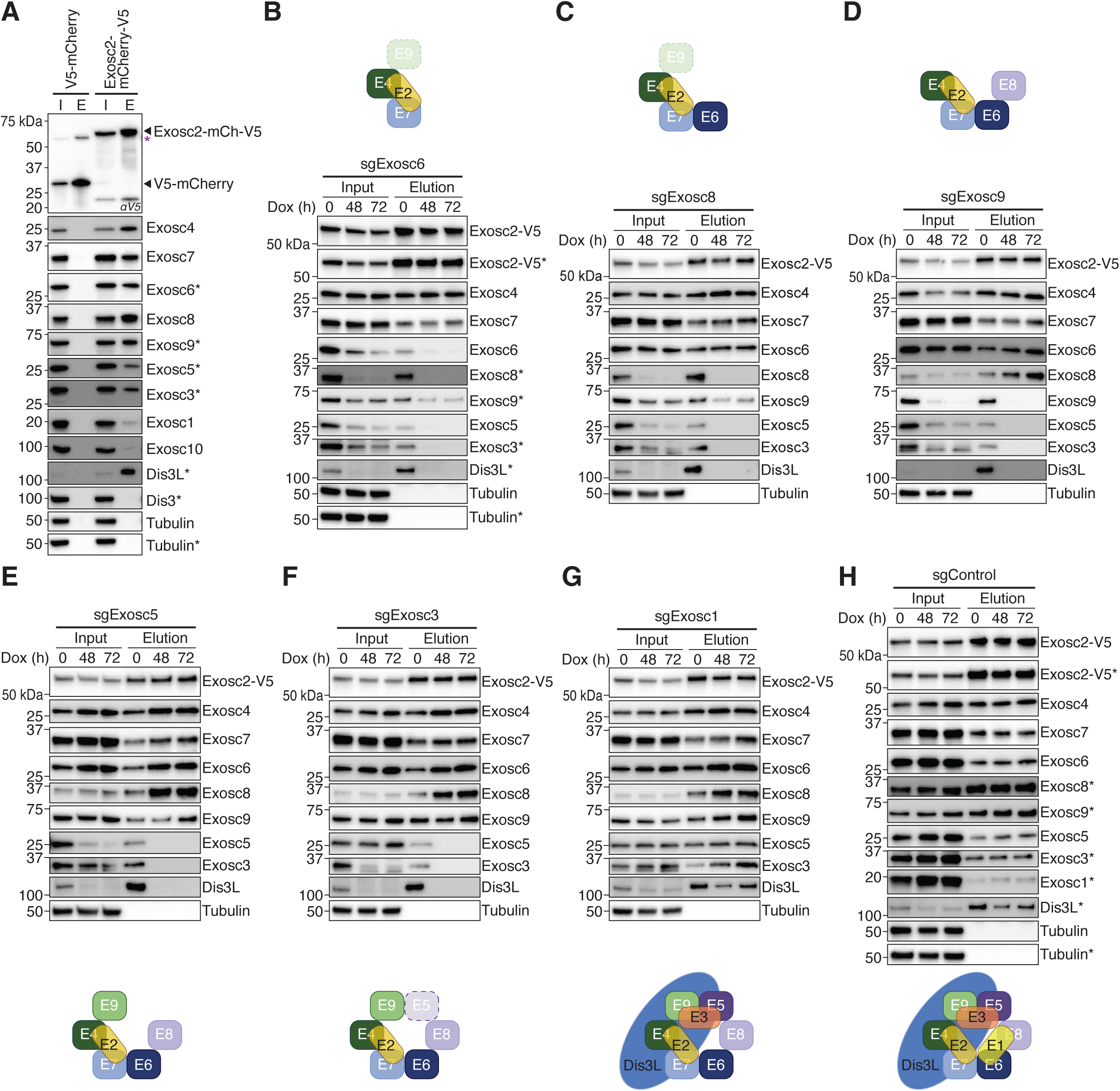
RNA exosome assembly intermediates form biochemically tractable subcomplexes. **(A)** V5-Immunoprecipitation (IP) was performed on lysate of mESCs expressing V5-mCherry-P2A-eBFP2 (control) or Exosc2-mCherry-V5-P2A-eBFP2. Tubulin serves as a loading control. Black asterisk (*) refers to matching membranes. Purple asterisk (*) refers to unprocessed V5-mCherry-P2A-eBFP2. **(B-H)** RNA exosome assembly intermediates were analyzed in lysate of mESCs expressing Exosc2-mCherry-V5-P2A-eBFP2, upon dual sgRNA-mediated depletion of Exosc6 **(B)**, Exosc8 **(C)**, Exosc9 **(D)**, Exosc5 **(E)**, Exosc3 **(F)**, Exosc1 **(G)**, or control Olfr10 **(H)** for the indicated time (in hours, h). *α*V5-IP was performed, and input and elution samples were analyzed by Western blotting. Schematics depict expected RNA exosome assembly intermediates. Asterisk (*) refers to matching membranes. See also **Figure S2**.

We interfered with RNA exosome assembly by depleting individual core components using iCas9 for 48 or 72 hours and examined complex formation by Exosc2-mCherry-V5 immunoprecipitation and Western blot analyses (Figure 3B-H). Indeed, this approach allowed us to isolate stage-specific RNA exosome assembly intermediates: Depletion of Exosc6 or Exosc8 resulted in the recovery of Exosc2/4/7 and Exosc2/4/7/6, respectively (Figure 3B and C). Both subcomplexes also retained Exosc9, consistent with the prediction that binding of Exosc9 to an immature RNA exosome core promotes resistance against orphan protein decay. In line with this assembly model, targeting Exosc9 by iCas9 also copurified the RNA exosome subcomplex Exosc2/4/7/6/8 (Figure 3D).

Strikingly, while Exosc5 was not destabilized upon the loss of Exosc3 (Figure S1N), immunoprecipitation of Exosc2-mCherry-V5 from Exosc3-iKO did not co-precipitate Exosc5 (Figure 3F), suggesting a weak or transient association of Exosc5 with the Exosc2/4/7/6/8/9 assembly intermediate. Provided that Exosc5 was co-purified together with Exosc2/4/7/6/8/9/3 in Exosc1-iKO (Figure 3G), this indicates that association of Exosc3 with the RNA exosome core is required for stable binding of Exosc5.

Finally, Exosc1 was not required for the stable association of Dis3L, consistent with the prediction that an Exosc1-deficient RNA exosome core can still form a catalytically competent complex (Figure 3G). Together with the fact that targeting negative control Olfr10 did not affect RNA exosome complex formation (ctrl-iKO; Figure 3H), and that the formation of subcomplexes was corroborated following the acute depletion of Exosc3 and Exosc6 using AID (Figure S2C and D), we concluded that depletion of RNA exosome core components stalls cellular RNA exosome assembly at specific stages that prevent orphan protein decay by forming biochemically stable subcomplexes.

### An immature core complex principally supports RNA exosome function and prevents apoptosis

To investigate if RNA exosome assembly intermediates retained activity, we focused on well-established RNA exosome targets, including promoter upstream transcripts (PROMPTs; Figure 4A), and rRNA precursors (i.e., pre-28S, pre-18S and pre-5.8S rRNA; Figure 4C)^19,20,21–23^. Using RT-qPCR, we measured the RNA abundance of four prototypic PROMPTs (proRpl23, proPou5f, proAnkhd1, and Fam120aos) following depletion of RNA exosome subunits or the Olfr10 control for 72 hours using iCas9 (Figure 4B). As expected, depletion of the cytoplasmic catalytic subunit Dis3L had no effect on PROMPT levels. In contrast, the loss of RNA exosome core components, as well as the nuclear catalytic subunits Exosc10 and Dis3, led to a significant upregulation of PROMPTs (p<0.05; two-way ANOVA and Holm-Šídák multiple comparison correction). Among the nuclear catalytic RNA exosome components, depletion of Exosc10 had a modest impact on PROMPT repression compared to Dis3, consistent with prior reports assigning a dominant role in PROMPT repression to Dis3^18,51^. Notably, depletion of components that joined the RNA exosome core at late assembly stages (i.e., Exosc5, Exosc3, and particularly Exosc1) caused consistently weaker derepression of PROMPTs compared to all other core components or Dis3 (Figure 4B). These findings, along with the independent validation of PROMPT upregulation as early as 2 hours following acute depletion of Exosc3 and Exosc6 using AID (Figure S3A and B), suggest that a minimal RNA exosome core structure lacking Exosc1 is partially compatible with Dis3-dependent decay of transcriptional byproducts.

**Figure 4.**
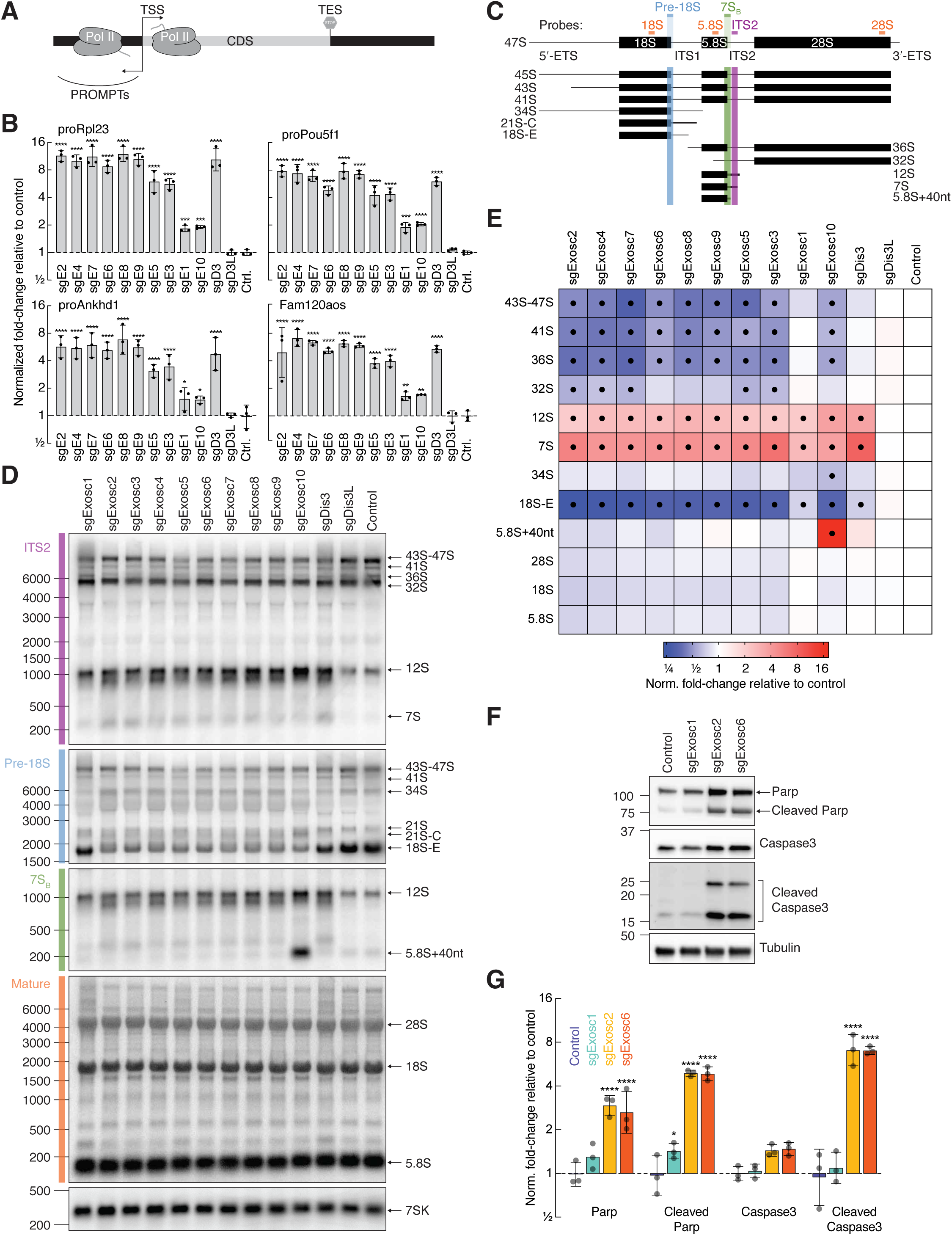
Functional assessment of RNA exosome assembly intermediates. **(A)** Schematic represents promoter upstream transcripts (PROMPTs), that are transcribed from the antisense strand upstream of active gene promoters. **(B)** Reverse transcription quantitative PCR (RT-qPCR) analysis of PROMPTs was performed in mESCs expressing dual sgRNAs targeting the indicated RNA exosome subunit upon treatment with Dox for 72h. Values were normalized to Gapdh mRNA and expressed relative to sg Olfr10 control. Bargraph reports mean ± SD of three independent replicates; individual measurements are indicated. Statistical significance was determined using two-way ANOVA with Holm-Šídák multiple comparison correction (* p_adj_ < 0.05, ** p_adj_ < 0.0021, *** p_adj_ < 0.0002, **** p_adj_ < 0.0001). **(C)** A schematic of rRNA processing intermediates is shown. Individual probes are indicated. **(D)** Northern blot analysis was performed on total RNA extracted from iCas9 mESCs expressing dual sgRNAs targeting the indicated RNA exosome subunits or Olfr10 control 72h after Dox treatment to detect the indicated rRNA processing intermediates. Probes used are shown in (C). **(E)** Average change in the abundance of rRNA processing intermediates as detected in (D), with rRNA intermediate abundance normalized to 7SK RNA and expressed relative to control (n = 3 biological replicates). Statistical significance was determined using two-way ANOVA with Holm-Šídák multiple comparison correction. Black dots indicate instances where p_adj_ < 0.05. **(F)** Western blot analysis was performed on lysate of iCas9 mESCs expressing dual sgRNAs targeting Olfr10 (control), Exosc1, Exosc2, and Exosc6, and treated with Dox for 72h. Tubulin serves as a loading control. **(G)** Quantification of (F), with protein abundance normalized to Tubulin and expressed relative to control (n = 3 biological replicates). Statistical significance was determined using two-way ANOVA with Holm-Šídák multiple comparison correction (* p_adj_ < 0.05, ** p_adj_ < 0.0021, *** p_adj_ < 0.0002, **** p_adj_ < 0.0001). **See also Figure S3, S4 and Table S4.**

To determine the impact of RNA exosome component depletion on rRNA processing, we employed Northern hybridization experiments using probes against specific rRNA biogenesis intermediates (Figure 4C). As expected, depletion of the cytoplasmic catalytic subunit Dis3L, similar to the Olfr10 control, did not significantly impact pre-rRNA processing (p>0.05; two-way ANOVA test and Holm-Šídák multiple comparison; Figure 4D and E). However, depletion of RNA exosome core components significantly affected the processing of Internal Transcribed Spacer (ITS) 1 (pre-18S, shown in blue, Figure 4C, D and E) and ITS2 (shown in purple, Figure 4C, D and E), as indicated by the downregulation of the rRNA processing intermediates 43S-47S, 41S, 36S, and 18S-E, and the upregulation of 7S and 12S (p<0.05; two-way ANOVA test and Holm-Šídák multiple comparison; Figure 4E). Furthermore, the loss of any RNA exosome core subunit led to the appearance of a non-discrete signal around the 18S-E precursor, a phenotype previously associated with the heterogenous population of 21S trimming intermediates, with 21S-C RNA representing the dominant species (Figure 4D)^22^. Notably, and as observed for PROMPTs (Figure 4B), depletion of Exosc1 resulted in a markedly weaker effect size compared to all others core subunits, suggesting that RNA exosome core structures lacking Exosc1 partially retained rRNA processing activity (Figure 4E).

Targeting of the nuclear catalytic subunit Dis3 using iCas9 primarily upregulated 7S and 12S rRNA precursors, underscoring its selective role in ITS2 processing. We observed only a slight reduction in the abundance of the 18S-E intermediate, consistent with the notion that this processing step occurs in the nucleolus while Dis3 is mainly localized in the nucleoplasm (Figure 4D and E)^52^.

Loss of Exosc10 resulted in a pronounced impairment of rRNA processing similar to that observed for the depletion of core subunits, but with the distinctive appearance of a longer isoform of the 5.8S precursor (Figure 4D and E). This finding aligns with prior observations that 7S intermediate is initially trimmed by Dis3 to approximately 40 nucleotides from the 3’ end of mature 5.8S rRNA, followed by further processing by Exosc10^22,53–55^.

To corroborate the rRNA processing defects observed upon the loss of RNA exosome core subunits, we subjected AID-Exosc3 and Exosc6-AID cell lines to an IAA treatment time-course (Figure S3C-F). Upregulation of 7S and 12S and downregulation of 18S-E precursors were evident after 2 hours of IAA treatment, while 43S-47S, 41S, and 36S intermediates became downregulated at later time points. These observations in the AID system align with the results obtained in the iCas9 system.

The weak functional phenotype observed in PROMPT repression and rRNA biogenesis upon Exosc1 depletion aligns with our previously noted mild effect on cell growth (Figure 1E). To further validate these results at the cellular level, we assessed apoptosis markers by Western blot analysis following the depletion of Exosc1 for 72 hours with iCas9 (Figure 4F)^56^. As expected, depletion of the early-assembled core components Exosc6 and Exosc2, but not Exosc1 or the negative control Olfr10, led to a substantial upregulation of cleaved Poly (ADP-ribose) Polymerase (PARP) and cleaved Caspase 3; Figure 4G).

In summary, we conclude that the terminal RNA exosome core component Exosc1 is partially dispensable for RNA exosome function at both the molecular and cellular levels. These findings highlight that the ordered assembly of core components establishes a functional hierarchy for the cell-essential repression of key RNA exosome targets.

### Exosc1 is dispensable for viability in mESCs

To determine whether mouse embryonic stem cells (mESCs) can tolerate a complete loss of Exosc1 protein, we generated three independent clonal knockout mES cell lines with frameshift mutations in the second exon of the Exosc1 gene using CRISPR/Cas9 genome engineering (Exosc1-KO; Figure S4A). The absence of Exosc1 protein was confirmed by Western blot analysis (Figure S4B) and targeted parallel reaction monitoring (PRM) mass spectrometry (Figures S4C-F). A competitive proliferation assay revealed moderate growth defects in clonal Exosc1-KO compared to wild-type (WT) mES cells (Figure S4G), similar to our observations with the iCas9 system (Figure 1E). To rule out off-target effects, we complemented Exosc1-KO cells with ectopic expression of N-or C-terminally V5-tagged Exosc1 (Figures S4H and I). Both V5-tagged Exosc1 constructs, but not the control mCherry-V5, rescued the proliferation defects (Figure S4J).

Functional characterization of the RNA exosome in Exosc1-KO clones showed moderate upregulation of PROMPTs (Figure S4K), which was fully rescued by re-expression of V5-tagged Exosc1 (Figure S4L). Analysis of rRNA processing in Exosc1-KO clones revealed a mild downregulation of the 18S-E precursor and weak upregulation of 7S and 12S precursors, consistent with results from inducible knockout experiments (Figure S4M and O). Re-expression of V5-tagged Exosc1 successfully restored normal rRNA processing (Figures S4N and P). The rRNA processing defects were more pronounced in the Exosc1-KO clones compared to Exosc1-depletion with iCas9 (Figure 4), likely due to terminal depletion effects in the clonal Exosc1 knockout.

Further analysis of global proteome alterations in Exosc1-KO mES clones using shotgun mass spectrometry revealed that among 5877 detected proteins, only 15 were significantly down- and eight upregulated, (p<0.05; limma and Benjamini Hochberg multiple comparison correction; Figure S4Q and Table S4). GO-term enrichment analyses for these differentially expressed proteins did not produce significant results. Reassuringly, Exosc10 stood out among the downregulated proteins (Figure S4Q and R), consistent with our observations using the iCas9 system (Figure 2), and confirmed by Western blot analyses (Figure S4S and T).

We concluded that long-term and complete depletion of Exosc1 has a minor impact on RNA exosome function, mES cell growth and global proteostasis. Hence, Exosc1 is dispensable for viability in mESCs.

## Discussion

Multiprotein complexes are ubiquitous, yet systematically understanding the role of individual subunits in their function, assembly and stability remains challenging, primarily due to technical limitations in protein depletion technologies^1–3^. We addressed these challenges by combining the scalability and ease of parallel gRNA-based gene targeting with doxycycline-inducible Cas9 (iCas9) expression, which accounts for indirect effects and the essential role of many protein complexes^35^. For gRNA design, we utilized the VBC score, which integrates gRNA activity, indel formation frequency, as well as amino acid composition and conservation of the targeted region to increase knockout efficiency^36^. By leveraging dual gRNA targeting, we further enhanced knockout penetrance within cell populations, mitigating the potential poor editing efficiency and in-frame repair of single sites. We benchmarked this approach against time- and labor-intense acute protein depletion strategies (i.e., auxin-inducible degron), demonstrating that the dual-gRNA iCas9 systems provides a robust and scalable method to systematically investigate direct proteostatic dependencies (Figure 1, S1).

By applying the dual-gRNA iCas9 system in mESCs to each of the twelve components of the RNA exosome (Figure 1C), we confirmed the essential cellular function of most RNA exosome subunits and observed specific patterns of exosome protein co-destabilization via the ubiquitin proteasome system (Figure 1D, 2A-F). These findings unveil the elaborate proteostatic control mechanisms governing an essential molecular machine that direct RNA processing and turnover of most coding and non-coding RNA species in eukaryotes^10^. In fact, most RNA exosome components, including all core components (i.e., Exosc1 to Exosc9), as well as the cytoplasmic Dis3L and the nuclear Exosc10, are subject to orphan protein decay (Figure 2A, B and S1G-R). These observations align with and significantly extend previous studies reporting selective orphan protein decay of RNA exosome components in *Trypanosoma brucei*, and human cells, suggesting that proteostatic control of the RNA exosome complex is evolutionarily conserved^13,16,19,29–34,49^.

Interestingly, despite their structural similarity, the nuclear Dis3, unlike its cytoplasmic paralog Dis3L, does not rely on association with the RNA exosome core for protein stability (Figure 2 and S1G-O). Both proteins depend on the PIN domain for RNA exosome association, yet Dis3 appears to escape proteostatic control that targets the cytoplasmic Dis3L^13,14^. Perhaps, Dis3 is specifically shielded prior to nuclear import or lacks destabilizing sequence features present in Dis3L. Alternatively, Dis3 may function independently of the RNA exosome core, as suggested by in vitro studies where yeast Rrp44 (Dis3) can bind and degrade RNA without the RNA exosome core^11,57,58^. However, our findings indicate that both Dis3 and the RNA exosome core are crucial for clearing PROMPTs or processing of 12S and 7S rRNA precursors, arguing against an independent role of Dis3 at least in these contexts (Figure 4B, D, and E).

Despite detailed information on the architecture and inter-subunit interactions emerging from seminal biochemical and structural work on reconstituted yeast and human RNA exosome complexes, how the complex is assembled remained a mystery^11,12,24–26,46,47,55,59,60^. Based on the selective proteolytic degradation of orphan RNA exosome components (Figure 2A, B and S1G-R), we propose a model for RNA exosome assembly that substantially differs from the previously proposed association of pre-formed dimers of barrel subunits in vitro, with cap components bridging the adjacent dimers^11^. In cells, assembly of the RNA exosome appears to be initiated by the formation of an interdependent heterotrimer comprised of the cap component Exosc2 and the barrel subunits Exosc4 and Exosc7. Formation of this trimer allows the sequential association of Exosc6, Exosc8, Exosc9, and Exosc5 to complete barrel formation, followed by the recruitment of Exosc3 and, lastly, Exosc1 to finalize the cap structure (Figure 2G).

While our model generally aligns with structural interaction data, it exhibits two notable exceptions during its cellular assembly. First, the stable association of Exosc9 with Exosc4 requires prior binding of Exosc8 to the Exo2/4/7/6 intermediate, implying a rearrangement that enables Exosc9 binding to Exosc4 (Figure 2A, B, 3B-D). Second, a reconfiguration must occur before Exosc1 recruitment and after Exosc3 binds to the Exosc2/4/7/6/8/9/5 complex (Figure 2A, B, 3F, G). Given the minimal direct contact between Exosc3 and Exosc1 in the human RNA exosome structure, Exosc3 may induce a cap-binding competent conformation of the barrel through direct interactions with Exosc9 and Exosc5, thereby facilitating Exosc1 binding to Exosc6 and Exosc8^11,46^. While the precise molecular nature of these rearrangements remains to be defined, they may involve changes in structural conformation exposing interaction sites, chaperon-assisted subunit incorporation, or the specific masking of confined destabilizing motifs only in the presence of specific assembly states.

The orphan protein decay of Exosc10 and Dis3L implies that catalytic subunit recruitment occurs at a late stage of RNA exosome assembly via distinct core subcomplexes (Figure 2, 3). Notably, Dis3 and Dis3L bind to the RNA exosome core through interactions with Exosc9, Exosc4 and Exosc7, which we predict form a complex before Exosc5, Exosc3, and Exosc1 recruitment^26,46–48^. Given that Dis3L protein stability depends on Exosc5 and Exosc3, but not Exosc1, stable Dis3L binding to the RNA exosome core must be masked prior to the second structural rearrangement induced by the recruitment of Exosc3 (Figure 2A, B). Based on their similar domain architecture, Dis3 may integrate into the RNA exosome core at a similar stage as Dis3L^13,14^. In contrast, Exosc10 stability depends on all RNA exosome core components, including Exosc1, indicating that its association relies on a completely assembled RNA exosome cap structure (Figure 2A, B). Overall, the presented assembly model provides a testable framework for future structural studies of experimentally tractable RNA exosome assembly intermediates (Figure 3).

The ordered assembly of the RNA exosome core complex establishes a functional hierarchy among its subunits (Figure 4). Early-assembled exosome component (i.e., Exosc2, Exosc4, Exosc7, Exosc6, Exosc8, and Exosc9) are equally essential for PROMPT repression, rRNA processing, and cell survival in mESCs. In contrast, depleting structural components that associate with the RNA exosome at late stages results in milder phenotypic consequences. For example, Exosc5 or Exosc3 depletion caused consistently weaker PROMPT derepression compared to other core components, perhaps because Dis3 retains the ability to associate with Exosc4, Exosc7 and Exosc9 in a core subcomplex that lacks Exosc5, Exosc3 and Exosc1^46–48^. However, both Exosc5 and Exosc3 are essential for rRNA processing, which largely depends on Exosc10, recruited to the RNA exosome core via Exosc6 and Exosc8, beyond Exosc1^46,47^. Strikingly, Exosc1, the final component in RNA exosome core assembly, is largely dispensable for PROMPT repression, rRNA processing, and mESC growth. This suggests that partial, low-affinity interactions of Dis3 and Exosc10 with the immature RNA exosome core may be sufficient to prevent apoptosis and sustain mESC viability, even upon complete, long-term depletion of Exosc1. In the absence of Exosc1, the RNA exosome may retain activity through interactions between the immature core and cofactors that provide substrate specificity, such as Mtr4 (Mtrex/Skiv2l2), C1d and Mphosph6, the latter two of which are only partially destabilized when Exosc1 is absent (Figure S4O-R)^24,46,61^.

The observed functional hierarchy among RNA exosome core components could explain the differential developmental roles of RNA exosome subunits in mouse embryos^28^. Exosc2 knockouts result in impaired uterine implantation at embryonic day (E) 3.5-4 and cannot be recovered at post-implantation stages, while Exosc1 knockouts progress further, forming an egg-cylinder but failing to initiate gastrulation or to develop beyond E5.5. Thus, while complete RNA exosome core assembly, including Exosc1, is dispensable for mESC viability and early embryonic development, it appears crucial for establishing multiple cell layers and coordinating body plan formation during gastrulation.

Overall, our findings establish a compelling framework for future investigations into the molecular basis, biological significance and biomedical relevance of proteostatic mechanisms that govern RNA exosome assembly, homeostasis, and function^31,62–71^.

## Author contributions

T.N. and S.L.A. conceived the project. T.N., with help of N.B., designed and performed the experiments. M.D., N.F. and V.H. contributed to cell line generation. K.S., G.K. and E.R. carried out proteomics experiments. R.W.K. and J.Z. designed dual gRNAs. N.P. performed computational analysis. T.N. and S.L.A. wrote the manuscript with input from all authors.

## Acknowledgments

We thank all members of the Ameres lab for technical support and helpful discussions; the IMP/IMBA/GMI and Max Perutz Labs core facilities for scientific support; Michael Schutzbier, Richard Imre, Gerhard Dürnberger and Otto Hudecz at IMP/IMBA/GMI Proteomics facility, Gerald Schmauß, Mario Nezhyba, Thomas Lendl, Marietta Weninger, Karin Aumayr at IMP Biooptics facility, Kitti Dora Csalyi, Endre Kiss and Thomas Sauer at Max Perutz Labs Biooptics facility for technical support; Maria Novatchkova (IMP/IMBA) for bioinformatics support; IMP/IMBA Molecular Biology service, members of Alexander Stark, Jan-Michael Peters and Andrea Pauli labs, Elif Karagöz, Gijs Versteeg and Egon Ogris labs for sharing reagents; and Sebastian Falk for discussions and comments on this manuscript. All proteomics analyses were performed using the Vienna BioCenter Core Facilities (VBCF) mass spectrometry instrument pool. T.N. was supported by a Boehringer Ingelheim Fonds PhD fellowship. Research in the S.L.A. lab is supported by the European Union (ERC, RiboTrace, CoG-866166), the Austrian Science Fund (FWF, 10.55776/F80 and 10.55776/DOC177), the Vienna Science and Technology Fund (WWTF, LS23-053), and the Austrian Academy of Sciences. For the purpose of Open Access, the authors have applied a CC BY license to any Author Accepted Manuscript (AAM) version arising from this submission.

## STAR Methods text Resource availability

### Lead contact

Further information and requests for resources and reagents should be directed to and will be fulfilled by the lead contact, Stefan L. Ameres.

### Materials availability

All unique reagents generated in this study are available from the lead contact without restriction.

### Data and code availability

- The mass spectrometry proteomics data have been deposited to the ProteomeXchange Consortium via the PRIDE partner repository with the identifier PXD061800. The processed proteomics data is available in this paper’s Tables S3 and S4.
- This paper does not report original code.
- Any additional information required to reanalyze the data reported in this paper is available from the lead contact upon request.

### Experimental model and subject details

#### mES and LentiX cell culture and chemical treatments

All presented experiments were performed in feeder-independent mouse embryonic stem cells (mESCs), clone AN3-12^72^. The mESCs were cultured in high-glucose DMEM (in house), supplemented with 13.5% fetal bovine serum (Sigma-Aldrich or Gibco), 1x Penicillin-Streptomycin (Sigma-Aldrich), 2mM L-Glutamine (Gibco) or 1x GlutaMax (Gibco), 1x MEM non-essential amino acid solution (Gibco), 1mM sodium pyruvate (Sigma-Aldrich), 50µM 2-mercaptoethanol and 20ng/ml LIF (in-house). Cells were kept at 37°C with 5% CO_2_ and passaged every second day.

For antibiotic selection in mESCs, Hygromycin (Roche) was used at 250µg/ml, BlasticidinS (Gibco) at 5µg/ml and Puromycin (Gibco) at 1µg/ml. For Cas9 induction, cells were cultured in Doxycycline-containing media (Dox, Sigma-Aldrich) at 500ng/ml for the indicated periods of time. For depletions in AID-tagged cell lines, indole-3-acetic acid sodium salt (IAA, Sigma-Aldrich) was used at 250µM final for indicated time-points. For proteasome inhibition Epoxomicin (Gentaur Molecular Products) was used at a final concentration of 1µM, and TAK-243 (ChemScence) was used for Uba1 inhibition at a final concentration of 2.5µM.

Lenti-X 293T cells (Clontech, 632180) were used only for lentivirus production. Cells were cultured in high-glucose DMEM (in house), supplemented with 10% fetal bovine serum (Sigma-Aldrich), 2mM L-Glutamine (Gibco) or 1x GlutaMax (Gibco), 1x MEM non-essential amino acid solution (Gibco), 1mM sodium pyruvate (Sigma-Aldrich), 50µM 2-mercaptoethanol. Cells were kept at 37°C with 5% CO_2_ and passaged every third day.

All cell lines were regularly tested to control for absence of mycoplasma contamination.

### Generation of Exosc1-KO cell line

To generate Exosc1-KO cells, gRNA targeting the second exon of Exosc1 was selected with a VBC score^36^. gRNA was cloned in pLenti-CRISPR-v2-GFP vector as described before^73^. Prior to transfection with gRNA containing plasmid, wild-type mESCs were stained with Hoechst33342 (Sigma-Aldrich) at 10µg/ml for 25min, followed by FACS-sorting (FACSAria III cell sorter, BD Biosciences) to obtain a haploid population. 5×10^5^ cells were reverse transfected with 2.5µg of Exosc1 gRNA-containing plasmid using Lipofectamine 2000 (Thermo Fisher Scientific) according to the manufacturer protocol. 48h post transfection GFP-positive cells were FACS-sorted and 2000 cells were seeded on a 15-cm dish. Single colonies were picked after 8 days and expanded. For genotyping, cells were lysed with DNA isolation buffer (10mM Tris-HCl pH 7.5, 10mM EDTA, 10mM NaCl, 0.5% N-Lauroylsarcosine, 1mg/ml Proteinase K) at 65°C for 2h followed by 95°C for 15 min. 1µl of the extracted genomic DNA was added to 20µl of PCR-reaction with primers flanking the edited loci. PCR-products were sanger sequenced and for selected clones the absence of Exosc1 was confirmed with Western blot and targeted PRM mass-spectrometry. Cells were stained with Hoechst33342 (Sigma-Aldrich) and diploid FACS-sorted prior to other experiments.

The gRNA sequence used to generate Exosc1-KO is indicated in Table S1.

### Generation of endogenously tagged AID-Exosc3 and Exosc6-linker-AID-V5 cell lines

A parental cell line expressing Tir1 E3-ligase was generated by inserting SFFV-Tir1-3xMyc-T2A-eYFP into Rosa26 loci by Flp-recombinase mediated cassette exchange (RMCE) into donor AN3-12 mESCs cells (GeneTrap Gt(Rosa26)Sor, Haplobank). To this end, 5×10^5^ cells were reverse co-transfected with a plasmid containing Tir1 flanked by FRT/F3 sites (pYR03_FRT-SFFV-OsTir1-3xMyc-T2A-eYFP-F3), plasmid expressing Flp (pCAG-Flp-GFP) at equimolar ratio, using Lipofectamine 2000 according to the manufacturer instruction. Seven days after transfection eYFP+ cells were FACS sorted and single clones were isolated as described in “Generation of Exosc1-KO mESCs cell line” section. Single clones were genotyped using integration site-specific PCR and Sanger sequencing. Cells were stained with Hoechst33342 (Sigma-Aldrich) and haploid FACS-sorted before proceeding with the endogenous tagging of Exosc3 and Exosc6.

For the endogenous tagging of Exosc3 and Exosc6, gRNAs targeting N-terminus of Exosc3 and C-terminus of were predicted with CHOPCHOP tool (v.3)^74^ and cloned into pX330-CMV-Cas9-hGem-P2A-mCherry, which was modified from pX330-CMV-Cas9-hGem (Addgene # 71707) by addition of P2A-mCherry. 5’- and 3’-homology arms were amplified from the genomic DNA and cloned along with AID-tag and V5-tag (for Exosc6) in pCR4-TOPO (Invitrogen) backbone. Next, haploid OsTir1-3xMyc-T2A-eYFP cells were reverse co-transfected (Lipofectamine 2000, Thermo Fisher Scientific) with an equimolar ratio of gRNA expressing plasmid and a homology template plasmid (5’ – homology arm – AID-tag – 3’-homology arm; pCR4-TOPO-AID-Exosc3-HT). For the endogenous C-terminal tagging of Exosc6 with linker-AID-V5-tag, haploid Tir1 cells were reverse co-transfected (Lipofectamine 2000, Thermo Fisher Scientific) with gRNA expressing plasmid and the homology template plasmid (5’ – homology arm – 3xGGGS linker – AID-tag – V5-tag – 3’-homology arm; pCR4-TOPO-Exosc6-L-AID-V5-HT) at 1:1 molar ratio. 48h post transfection mCherry+ cells were FACS sorted and single clones were isolated as described in “Generation of Exosc1-KO mESCs cell line” section. Successful protein tagging was validated with the integration site-specific PCR, Sanger sequencing and Western blot. Note, that AID-tagging of Exosc3 resulted in a modest but measurable reduction in Exosc3 protein levels compared to the parental cell line (Figure S1E), in line with a previous report of Exosc3 AID-tagging^19^.

Sequences of gRNAs used for endogenous AID-tagging of Exosc3 and Exosc6 are indicated in Table S1.

### Generation and validation of iCas9v1 cell line

As a parental cell line for iCas9v1 generation we used previously generated in the lab for other purposes OsTir1-3xMyc-T2A-eYFP expressing Xrn1-L-AID-V5 cells. 300k cells were seeded on a 6-well plate and sequentially transduced with pRRL-SFFV-rtTA3-IRES-EcoRec-PGK-Hygro and pRRL-TRE3G-Cas9-P2A-eBFP2 and selected with Hygromycin B. Next, Doxycycline (DOX, Sigma-Aldrich) was added to induce eBFP2 expression and eBFP2+ cells were FACS sorted followed by single cell clone isolation as described in “Generation of Exosc1-KO mESCs cell line” section. To assess the function of Cas9 and tightness of TRE3G promoter in the selected clones, cells were transduced with the construct expressing gRNA eYFP and Cas9 expression was induced with DOX (Sigma-Aldrich). Competitive proliferation assays were set-up to monitor the reduction of eYFP signal upon its knockout. All measurements were performed with flow cytometry on iQue Screener PLUS (Intellicyt) or LSRFortessa (BD Biosciences).

### Generation of iCas9v2 cell line

For the generation of iCas9v2, we inserted all-in-one cassette, containing SFFV-rtTA3-IRES-HygroR-insulator-insulator-iRFP713-P2A-Cas9-TRE3G, into the expression-stable locus on Chr15^75^ with Flp-recombinase mediated cassette exchange (RMCE). 5×10^5^ cells were reverse co-transfected with a plasmid containing the cassette flanked by FRT/F3 sites (pYR03-FRT-Cbh-rTetR-IH-ins2-iRFP713-P2A-Cas9-TRE3G-ins2-F3), and a plasmid expressing Flp (pCAG-Flp-GFP) at equimolar ratio, using Lipofectamine 2000 according to the manufacturer instruction. Seven days after transfection cells were FACS-sorted and single clones were isolated as described in “Generation of Exosc1-KO mESCs cell line” section. Single clones were genotyped using integration site-specific PCR and Sanger sequencing. To assess the function of Cas9 and tightness of TRE3G promoter, clones were transduced with the construct expressing gRNA against CD9 followed by Cas9 induction and CD9-immunostaining.

### Surface marker immunostaining

Cells were pelleted (200g, 3 min, 4°C) and stained with eFluor450-conjugated anti-CD9 antibody (1:500, Thermo) in FACS buffer (1x PBS, 5% FBS) for 10 min at 4°C. Cells were washed once with FACS buffer, spun down, resuspended in FACS buffer, and analysed with flow cytometry using LSRFortessa (BD Biosciences) and Penteon (Novocyte).

### Competitive proliferation assays

Proliferation analyses of inducible knockouts were performed as follows: dual gRNA mCherry+ iCas9 cells were mixed with a parental iCas9 cell line, DOX was added to induce Cas9 expression and a fraction of mCherry+ cells was monitored over time with flow cytometry.

For growth competition experiments in AID-tagged cell lines (in Tir1-T2A-eYFP background), AID-Exosc3 and Exosc6-AID cells were co-cultured with wild-type cells in the absence/presence of indol-3-acetic acid (IAA, Sigma-Aldrich). Percentage of eYFP+ cells was followed over time with flow cytometry.

To address proliferation defects of monoclonal Exosc1-KO and rescue cell lines, respective cell lines were co-cultured with OsTir1-3xMyc-T2A-eYFP cells and percentage of eYFP-cells was derived from flow cytometry measurements.

All measurements were performed with iQue Screener PLUS (Intellicyt) or LSRFortessa (BD Biosciences). Data analysis was done with FlowJo (v.10.9.0) software.

### Design and cloning of dual sgRNAs

The top performing gRNA from the Vienna gRNA^36^ library was used to predict pairing gRNAs. All gRNAs targeting murine exosome genes were filtered stringently for off-targets, and polyA and restriction-site containing sequences were removed. Then, relative cutting positions to the original guide sequences were calculated and filtered for a window of +/- 50 – 1000 bp, and further filtered for out-of-frame cut site distances. Resulting matching gRNAs were ranked by VBC score and the top guide was selected.

Dual gRNAs were ordered as oligonucleotides (see Table S1), annealed, and cloned into pDual-hU6-sgRNA-mU6-sgRNA-EF1α-mCherry-P2A-PuroR expression vector using BbsI-HF (NEB) and Esp3I (NEB) restriction enzymes.

### Cloning of constructs for exogenous expression

Expression vectors for the ectopic expression of Exosc1 and Exosc2 were based on pRRL backbone (Addgene #145840). pRRL vector was modified to contain CMV promoter and BlasticidinS resistance cassette (only for Exosc2).

Full-length cDNA for Exosc1 (NM_025644.4) and Exosc2 (NM_144886.3) was prepared with reverse-transcription from total RNA of wild-type mESCs, followed by a PCR amplification of the region containing an open reading frame. Fragments for mCherry, eBFP2 and mScarlet3 were separately amplified and linker sequence (3xGGGS), V5-tag and SPOT-tag were introduced with the primers.

All described constructs were assembled with NEBuilder HiFi DNA Assembly Master Mix (NEB) from the respective fragments and verified by Sanger sequencing. List of plasmids is in Key resource table.

### Lentivirus production and infections

For lentiviral production 4×10^5^ LentiX cells were plated on a 6-well plate and transfected on the next day. To prepare a transfection mixture 2300ng of donor plasmid (i.e. expressing dual sgRNAs) were mixed with 1140ng pCMVR8.74 (Addgene #22036), 670ng pMD2.G (Addgene #12259), 12µl polyethylenimine (PEI, Polysciences) in 200µl OPTI-MEM (Gibco). Transfection mix was vortexed, incubated for 25 min at room temperature and added dropwise to the cells. Media was changed 4-10 h after transfection and lentiviruses were harvested 24 h later. Viral supernatant was filtered through 0.45µm filter and used for infection of mESCs in the presence of 4µg/ml polybrene (Merck). List of plasmids used for lentiviral transduction can be found in Key resource table.

### Western blotting

Whole cell protein extracts were prepared by resuspending cells in the ice-cold Western blot lysis buffer (50mM Tris-HCl pH 7.5, 150mM NaCl, 1% Triton X-100, 0.5% Tergitol, 0.1% SDS, 0.5mM EDTA) supplemented with 1x fresh HALT protease inhibitors (Thermo). Lysates were incubated on ice for 30 minutes and spun down at 20000g, 4°C for 20 minutes. Supernatant, containing total protein lysate, was collected and its concentration was determined with Bio-Rad Protein Assay Dye Reagent (Bio-Rad) according to the manufacturer’s instructions. 50-150µg of total protein were mixed with 6X loading buffer (375mM Tris-HCl pH 6.8, 12% SDS, 60% glycerol, 6% 2-mercaptoethanol, bromphenol blue) and denatured at 95°C for 5 min. Samples were separated either on self-made acrylamide gels (5% stacking, 12% separating) or on 4-20% pre-cast gradient gels (Bio-Rad) and transferred to Immobilon-P PVDF membrane (EMD Millipore) at 100V for 1h using a wet-chamber system. Membranes were blocked with 3% non-fat milk in TBS + 0.05% Tween-20 (TBST) for 1h at room temperature and incubated with primary antibodies at 4°C overnight. Next day, membranes were washed 5 x 5 min with TBST and HRP-secondary antibody were added for 1h at room temperature followed by 5 x 5 min TBST washes. Proteins were detected using Clarity Western ECL Substrate (Bio-Rad) with ChemiDoc Imaging System (Bio-Rad) using ImageLab V5.1.1 (Bio-Rad). For the membranes that were probed with several different antibodies, membranes were incubated in the Restore Western blot stripping buffer (Thermo) at room temperature for 20 min followed by 3 x 5 min TBST washes, re-blocking and incubation with primary/secondary antibody.

The following antibodies were used (also see the key resource table): rabbit anti-Exosc1 (1:5000, Bethyl), rabbit anti-Exosc2 (1:2000, Bethyl), rabbit anti-Exosc2 (1:5000, Proteintech), mouse anti-Exosc2 (1:3000, Proteintech), rabbit anti-Exosc3 (1:1000, Novus), rabbit anti-Exosc3 (1:5000, Proteintech), rabbit anti-Exosc4 (1:5000, Proteintech), mouse anti-Exosc4 (1:3000, Santa Cruz), rabbit anti-Exosc5 (1:2000, Proteintech), rabbit anti-Exosc6 (1:5000, Thermo), mouse anti-Exosc6 (1:1000, Santa Cruz), mouse anti-Exosc7 (1:200, Santa Cruz), rabbit anti-Exosc7 (1:10000, Proteintech), rabbit anti-Exosc8 (1:3000, Proteintech), rabbit anti-Exosc9 (1:3000, Proteintech), mouse anti-Exosc9 (1:3000, Proteintech), mouse anti-Exosc9 (1:1000, Santa Cruz), mouse anti-Exosc10 (1:1000, Santa Cruz), rabbit anti-Dis3 (1:3000, Proteintech), mouse anti-Dis3L (1:100, Santa Cruz), rabbit anti-C1d (1:1000, Proteintech), rabbit anti-Mphosph6 (1:1000, Proteintech), rabbit anti-Parp (1:5000, Cell Signalling), rabbit anti-Caspase3 (1:3000, Cell Signalling), rabbit anti-Cleaved Caspase3 (1:2000, Cell Signalling), rabbit anti-Actin (1:5000, Sigma-Aldrich), mouse anti-Tubulin (1:20000, Proteintech), mouse anti-Hsp70 (1:1000, Abcam), mouse anti-V5 (1:5000, Thermo), mouse anti-SPOT (1:5000, Proteintech), HRP-conjugated anti-mouse (1:10000, Thermo) and HRP-conjugated anti-rabbit (1:10000, Thermo).

Western blot quantifications were done with ImageJ software (v. 2.14.0) taking into account the linear range of chemiluminescence signal^76^ as judged from the dilution series (100%, 50%, 25%, 10%). Often the band was not visible in the lowest 10% dilution sample, thus we fixed 10% as the detection limit of our Western blot analyses and reported the maximum depletion efficiency of being less or equal 10%. For all the quantifications the signal of a protein of interest was normalized to the loading control (Hsp70, Actin or Tubulin) and presented relative to control samples.

### Immunoprecipitation experiments

Cells were resuspended in ice-cold IP lysis buffer (50mM Tris-HCl pH 7.5, 150mM NaCl, 1% Tergitol, 0.5mM EDTA) supplemented with 1x fresh HALT protease inhibitors (Thermo). Lysates were incubated on ice for 30 min, sonicated (40% power, 3 cycles, 10 sec, Bandelin Sonoplus HD2070) and centrifuged at 20000g, 4°C for 20 minutes. Supernatant, containing total protein lysate, was collected and its concentration was determined with Bio-Rad Protein Assay Dye Reagent (Bio-Rad) according to the manufacturer’s instructions. 1mg of total protein was mixed with either 25µl of V5-Trap Magnetic Agarose (Proteintech) or 20µl of SPOT-Trap Magnetic Agarose (Proteintech) and incubated at 4°C for 4h with end-to-end mixing. Beads were washed 4 times with IP lysis buffer and proteins were eluted with 2X loading buffer at 95°C for 10 min. Input and elution samples were resolved on 4-20% pre-cast gels (Bio-Rad) and Western blotting was performed as described above.

### RNA isolation and RT-qPCR

Total RNA was isolated using 1ml of Trizol (Thermo) according to the provider’s protocol. Isolated RNA was subjected to Turbo DNAse (Thermo) treatment following the manufacturer’s instructions. 2µg of total RNA were used for cDNA preparation with Superscript III reverse transcriptase (Thermo) and a mix of 20µM oligo dT20 and 50ng/µl random primers (Thermo). qPCR was performed with GoTaq qPCR Master Mix (Promega) using CFX 384 Touch (Bio-Rad) Real-Time PCR with Bio-Rad CFX Manager (v.1.0) software. In all cases melting curves were assessed and selected products were run on an agarose gel to ensure that only a specific product was amplified. qPCR primers are listed in Table S2.

qPCR signal for targets of interest was quantified using 2^-ΔΔCt^ method^77^, normalized to an internal control (Gapdh mRNA) and presented as a fold-change relative to the control sample.

### Northern blotting

8µg of total RNA were resolved on 1.2% agarose gel containing 1x MOPS buffer (40mM MOPS, 10mM sodium acetate, 1mM EDTA; pH of the whole buffer was adjusted to 7.2) and 2% formaldehyde. Before loading on the gel, RNA was mixed with 2X loading buffer (2x MOPS buffer, 9% formaldehyde, 0.3x gel loading buffer II (Thermo), 20 µg/ml ethidium bromide) incubated at 75°C for 10 min and quickly chilled on ice. After the run, RNA was transferred onto N+ Hybond membrane (Thermo) by an overnight capillary transfer. The membrane was UV cross-linked (Auto mode of UV Stratalinker 2400, twice) and pre-hybridized with a Church Buffer (0.5M Na_2_HPO_4_/NaH_2_PO_4_ pH 7.2, 1mM EDTA, 7% SDS) at 65°C for 1h followed by the addition of the radiolabeled probe and overnight incubation at 37°C unless indicated otherwise. Next day, the membrane was washed 3 x 15 min with 1X SSC (150mM NaCl, 15mM sodium citrate; pH of the whole buffer was adjusted to 7.0) + 0.1% SDS buffer and radiation was recorder with a phosphor screen (GE Healthcare) for 24-48h. Imaging was done on Amersham Typhoon phosphor imager (Cytiva). To proceed with the next probe, membranes were incubated in 0.1% SDS at 95°C for 15 min, followed by pre-hybridization and incubation with a new radiolabeled probe. Probes were used in the following order: ITS2_probe (incubation at 42°C) => Pre-18S_probe => 7S_B_ probe => 7SK probe => mix of 5.8S, 18S, 28S probes. Northern blot probes are listed in Table S2.

Northern blots were quantified using ImageQuant TL (v.10.2) software. Intensities of bands of interest were normalized to a loading control (7SK RNA) and presented relative to control samples.

### Peptide preparation for the mass-spectrometry

For Exosome inducible knockout experiments cells were seed on a 6-well plate and treated for 72h with Dox. For monoclonal Exosc1-KO experiments, wild-type and Exosc1-KO clones were seed on a 6-well plate 24h before the harvest. Next, cells were washed twice with 1X PBS, harvested with a cell scraper and spun down at 200g for 3 min. Cell pellet was frozen in liquid nitrogen and stored at −80°C. Peptides for mass-spectrometry analysis were prepared with iST kit (Preomics) according to the manufacturer’s instructions with slight modifications. Frozen cell pellets were incubated with 50µl of LYSE at 95°C, 10min, 1000 rpm. Lysates were sonicated for 30 seconds (10 cycles, 50% amplitude, UP100H, Hielscher) and total protein concentration was determined with Micro BCA kit (Thermo). 100µg of total protein were mixed with 50µl DIGEST and incubated overnight at 37°C. Next day, digestion was quenched with 100µl of STOP solution and the sample was applied to a cartridge followed by WASH1 and WASH2. Cleaned peptides were eluted twice with 100µl of ELUTE and placed in the SpeedVac machine (Concentrator plus, Eppendorf, 45°C, V-AQ mode) until all the liquid evaporated. Sample was resuspended in 100µl of 0.1% trifluoroacetic acid (TFA) and sonicated in an ultrasonic bath (Bandelin Sonorex super RK31) for 5 min to solubilize the peptides.

Final peptide amount was determined by separation of an aliquot of the sample on LC-UV system equipped by monolith column (Thermo) based on the peak area of 100ng of HeLa protein digest standard (Thermo). Peptide solution was frozen at - 80°C before further processing.

### Shotgun LC-MS/MS analysis

The nano-HPLC system UltiMate 3000 RSLC was coupled to an Orbitrap Exploris 480 mass spectrometer, equipped with a FAIMS pro interface and a Nanospray Flex ion source (all parts Thermo). Peptides were loaded onto a trap column (PepMap C18, 5mm × 300μm ID, 5μm particles, 100Å pore size, Thermo). The trap column was switched in line with the analytical column (PepMap C18, 75μm ID × 50cm, 2μm, 100Å, Thermo). The analytical column was connected to PepSep sprayer 1 (Bruker) equipped with a 10μm ID fused silica electrospray emitter with an integrated liquid junction (Bruker, Part#1893527). Electrospray voltage was set to 2.4kV. Peptides were eluted using a flow rate of 230nl/min, starting with the mobile phases 98% A (0.1% formic acid in water) and 2% B (80% acetonitrile, 0.08% formic acid) and linearly increasing to 35% B over the next 180 min, followed by a gradient to 95% B in 5 min, staying there for 5 min. Column equilibration was done at 30°C with 2% B for 25min for U3000.

The Orbitrap Exploris 480 mass spectrometer was operated in data-dependent mode, performing a full scan (m/z range 350-1200, resolution 60,000, normalized AGC target 100%) at 3 different compensation voltages (CV −45, −60, −75), followed each by MS/MS scans of the most abundant ions for a cycle time of 0.8. MS/MS spectra were acquired using a collision energy of 30, isolation width of 1.0 m/z, resolution of 15.000, max fill time 100ms, normalized AGC target of 100 % and intensity threshold of 2.5×10^4^. Precursor ions selected for fragmentation (include charge state 2-6) were excluded for 40s. The monoisotopic precursor selection (MIPS) filter and exclude isotopes feature were enabled.

### Proteomics data analysis

Raw MS data was loaded into Proteome Discoverer (v. 2.5.0.400). All MS/MS spectra were searched using MS Amanda (v2.0.0.19924)^78^. Trypsin was specified as a proteolytic enzyme cleaving after lysine and arginine without proline restriction, allowing for up to 2 missed cleavages. Mass tolerances were set to ±10ppm at the precursor and fragment mass level. Peptide and protein identification was performed in two steps. An initial search was performed against the reference database (UP000000589, 21,962 sequences; 11,728,099 residues), with common contaminants appended. Here, carbamidomethylation of cysteine was searched as fixed modification, whereas oxidation of methionine, deamidation of asparagine and glutamine and glutamine to pyro-glutamate conversion at peptide N-termini were defined as variable modifications. Results were filtered for a minimum peptide length of 7 amino acids and 1% FDR at the peptide spectrum match (PSM) and the protein level using the Percolator algorithm^79^ integrated in Proteome Discoverer. Additionally, an Amanda score of at least 150 was required.

A sub-database of proteins identified in this search was generated and used for a second search, where the RAW-files were analysed using the same settings as above plus considering additional variable modifications: phosphorylation of serines, threonines and tyrosines, acetylation on protein N-Terminus. The localization of the post-translational modification sites within the peptides was performed with the tool ptmRS, based on the tool phosphoRS^80^. Identifications were filtered using the criteria described above, including an additional minimum PSM-count of 2 per protein in at least one sample. The identifications were subjected to label-free quantification using IMP-apQuant^81^. Proteins were quantified by summing unique and razor peptides and applying intensity-based absolute quantification (iBAQ)^37^ with subsequent normalization based on the MaxLFQ algorithm^82^. Identified proteins were filtered to contain at least 3 quantified peptide groups. Statistical significance of differentially expressed proteins was determined using limma^83^. GO-term enrichment analysis was done with clusterProfiler R package^84^.

All proteomics samples were measured in biological triplicates, except for inducible knockout of Exosc5 (sgExosc5; Figure 2B) where two replicates were analysed.

Results of the proteome comparison between inducible knockouts of Exosc1-Exosc10, Dis3, Dis3L, Olfr10 and Olfr1 are listed in Table S3.

Results of the proteome comparison between wild-type and Exosc1-KO are listed in Table S4.

### Targeted mass spectrometry via parallel reaction monitoring (PRM)

Vanquish Neo UHPLC-System or UltiMate 3000 RSLC coupled to the Orbitrap Exploris 480 mass spectrometer, equipped with an Easy spray Source (Thermo) were used. Peptides were loaded onto a trap column (PepMap C18, 5mm × 300μm ID, 5μm particles, 100Å pore size, Thermo) by using 0.1% TFA. The trap column was switched in line with the analytical column (PepMap C18, 500mm × 75μm ID, 2μm, 100Å, Thermo). The analytical column was connected to PepSep sprayer 1 (Bruker) equipped with a 10μm ID fused silica electrospray emitter with an integrated liquid junction (Bruker, Part#1893527). Electrospray voltage was set to 2.1kV. Peptides were eluted at 230nl/min using a binary 90-minute gradient within the complete 135-minute LC-MS/MS run, including column equilibration.

The gradient starts with the mobile phases: 98% A (water/formic acid, 99.9/0.1, v/v) and 2% B (water/acetonitrile/formic acid, 19.92/80/0.08, v/v/v), increases to 35% B over the next 90min, followed by a gradient in 5min to 90% B. Column equilibration was done with 2% B by 3 analytical column volumes in case of Neo Vanquish or for 25min in case of U3000 at 30°C.

The Orbitrap Exploris 480 mass spectrometer was operated by a mixed MS method which consisted of one full scan (*m/z* range 380-1,500; resolution 15,000; target value 100%) followed by the PRM of targeted peptides from an inclusion list (isolation window 0.8 *m/z*; normalized collision energy (NCE) 32; resolution 30,000, AGC target 200%). The maximum injection time was set to 800ms.

A scheduled PRM method (sPRM) development, data processing and manual evaluation of results were performed in Skyline (v. 22.2.0.351)^85^. Spectra of unique peptides of the proteins of interest (Exosc1, Exosc2, Exosc3, Dis3L and C1d) and normalization controls (Cct8, Eef2, Atp5f1b, Ncl, Pkm, L1td1) were recorded. Each peptide used for relative quantification of a protein of interest had to have at least 3 fragment ions.

For each protein of interest and normalization control, measured peptide areas were summed up. Obtained value for each protein of interest was divided by the value corresponding to the individual normalization control and presented relative to the respective control sample (wild-type for Exosc1-KO and sgOlfr1 for Exosome inducible knockout experiments). Median value (out of all normalization controls) was used as a relative estimation of the protein abundance.

For Figure 2B, PRM was used to determine relative protein abundance of Exosc1, Exosc2, Exosc3 and Dis3L. In sgExosc2 (two out of three replicates), sgExosc7 and sgExosc8 samples no signal for Dis3L peptides was detected. Consequently, the Dis3L abundance was set to the maximum observed co-depletion.

For Figure S4R, PRM was used to determine relative protein abundance of Exosc1, Exosc3, Dis3L and C1d.

Results of the PRM measurements in inducible knockouts of Exosc1-Exosc10, Dis3, Dis3L, Olfr10 and Olfr1 are listed in Table S3.

Results of the PRM measurements in wild-type and Exosc1-KO are listed in Table S4.

### Quantification and statistical analysis

Quantification methodologies of Western blots, northern blots, qPCRs and mass-spectrometry are described in the respective sections. Statistical analyses are indicated in the figure legends.

**Table S3. Proteome comparison of inducible knockouts of Exosc1-Exosc10, Dis3, Dis3L, Olfr10 and Olfr1**, related to Figure 2 and S1.

**Table S4. Proteome comparison of Exosc1-KO and wild-type cells,** related to Figure S4.

## Key resources table

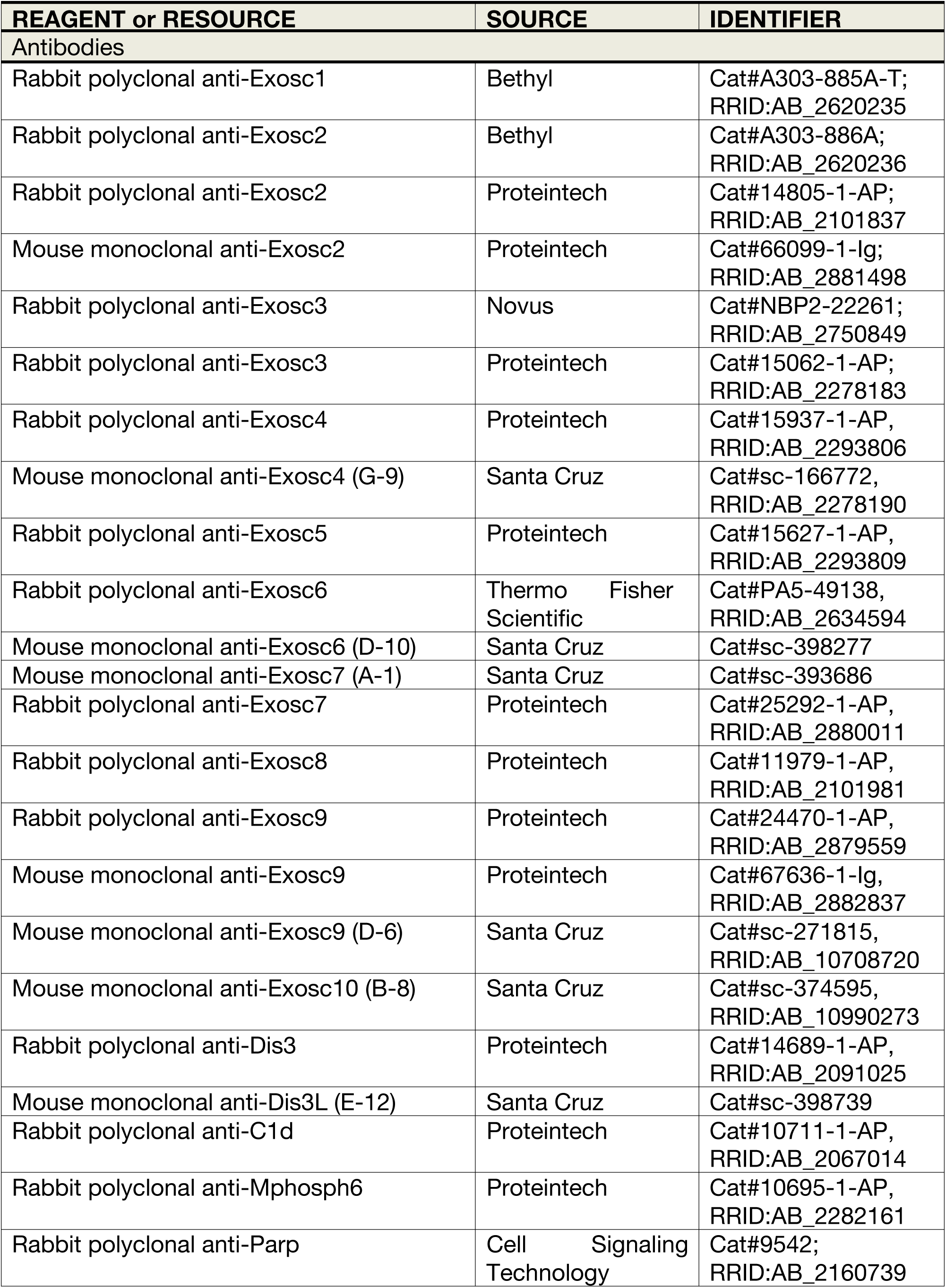
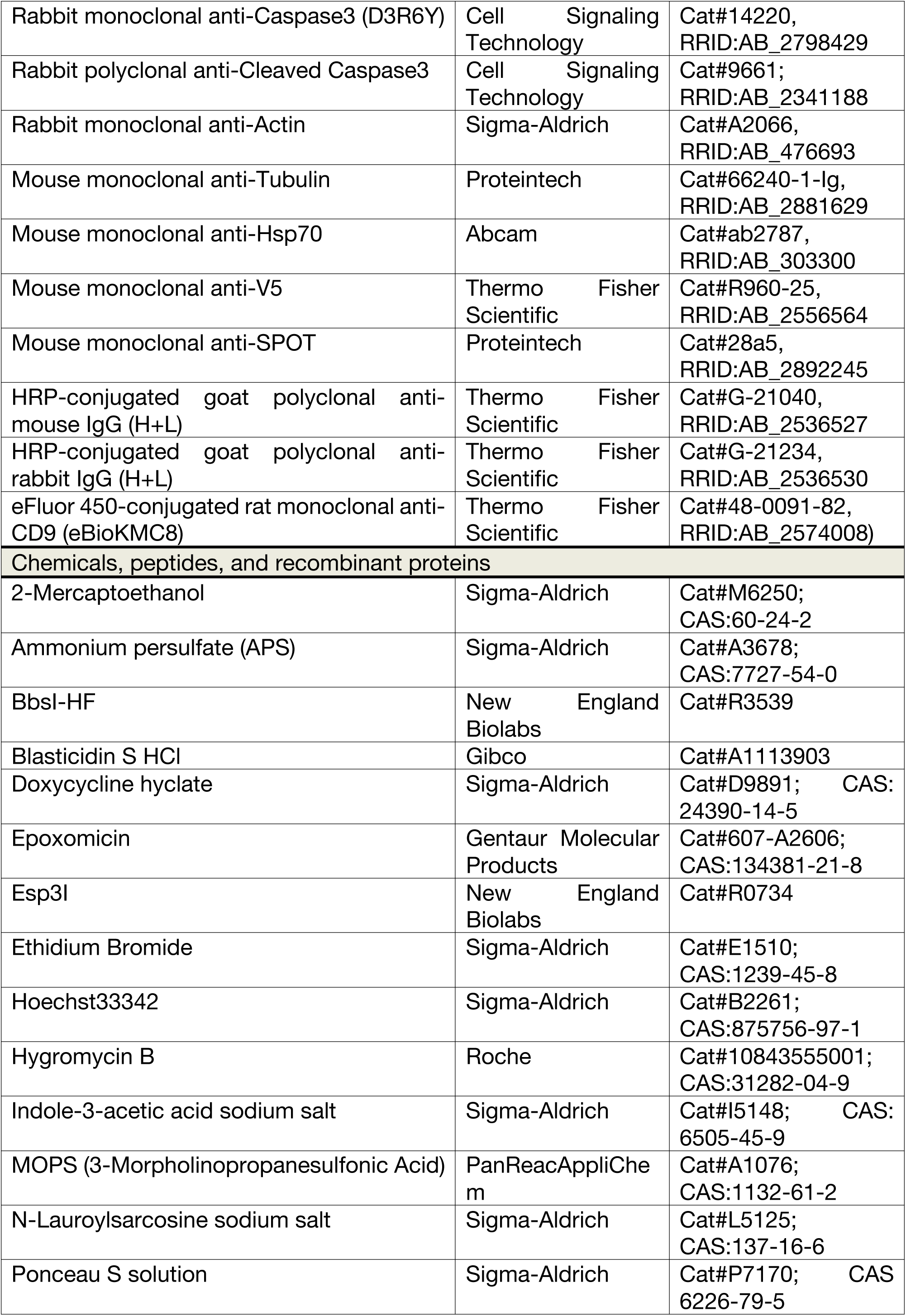
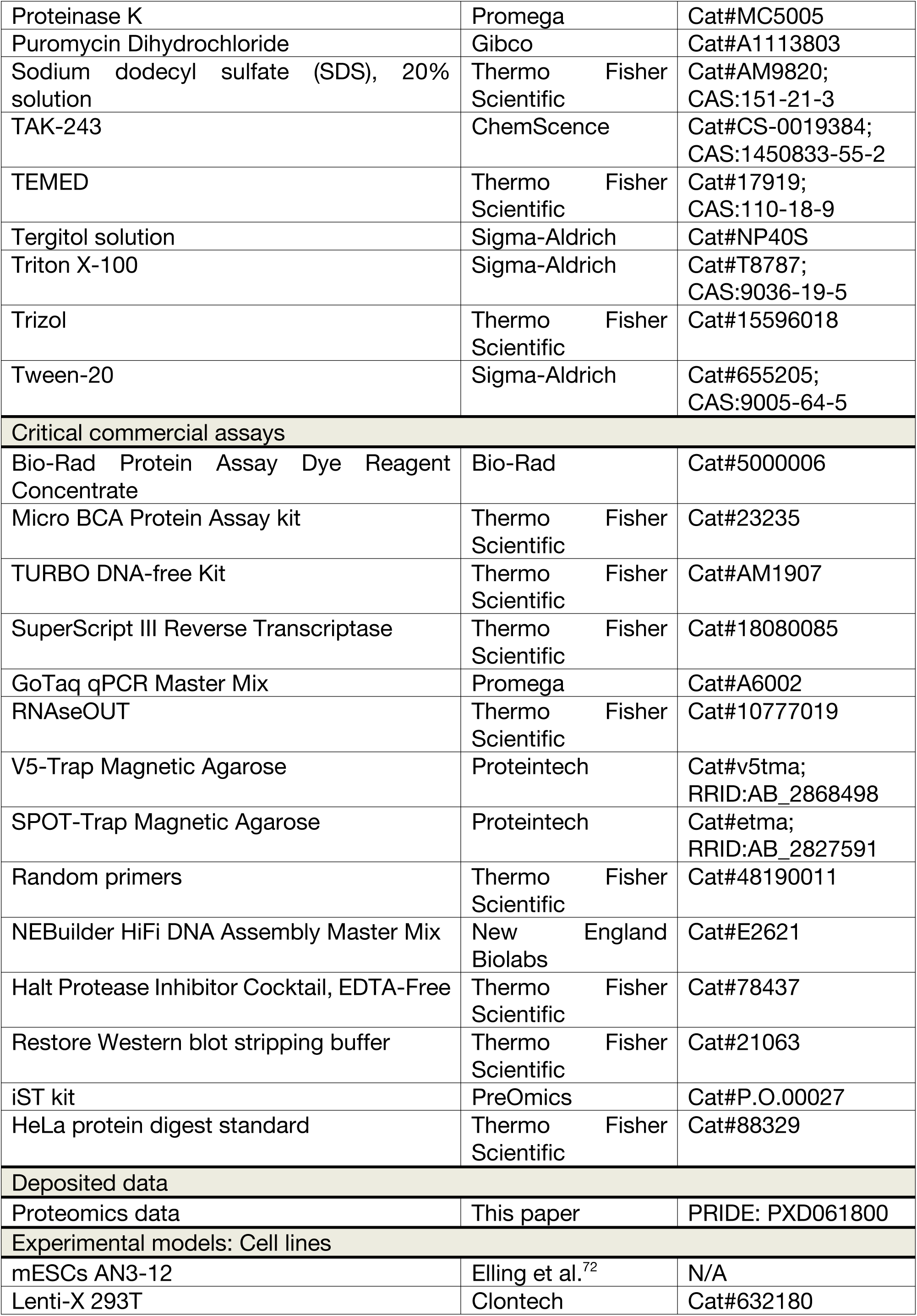
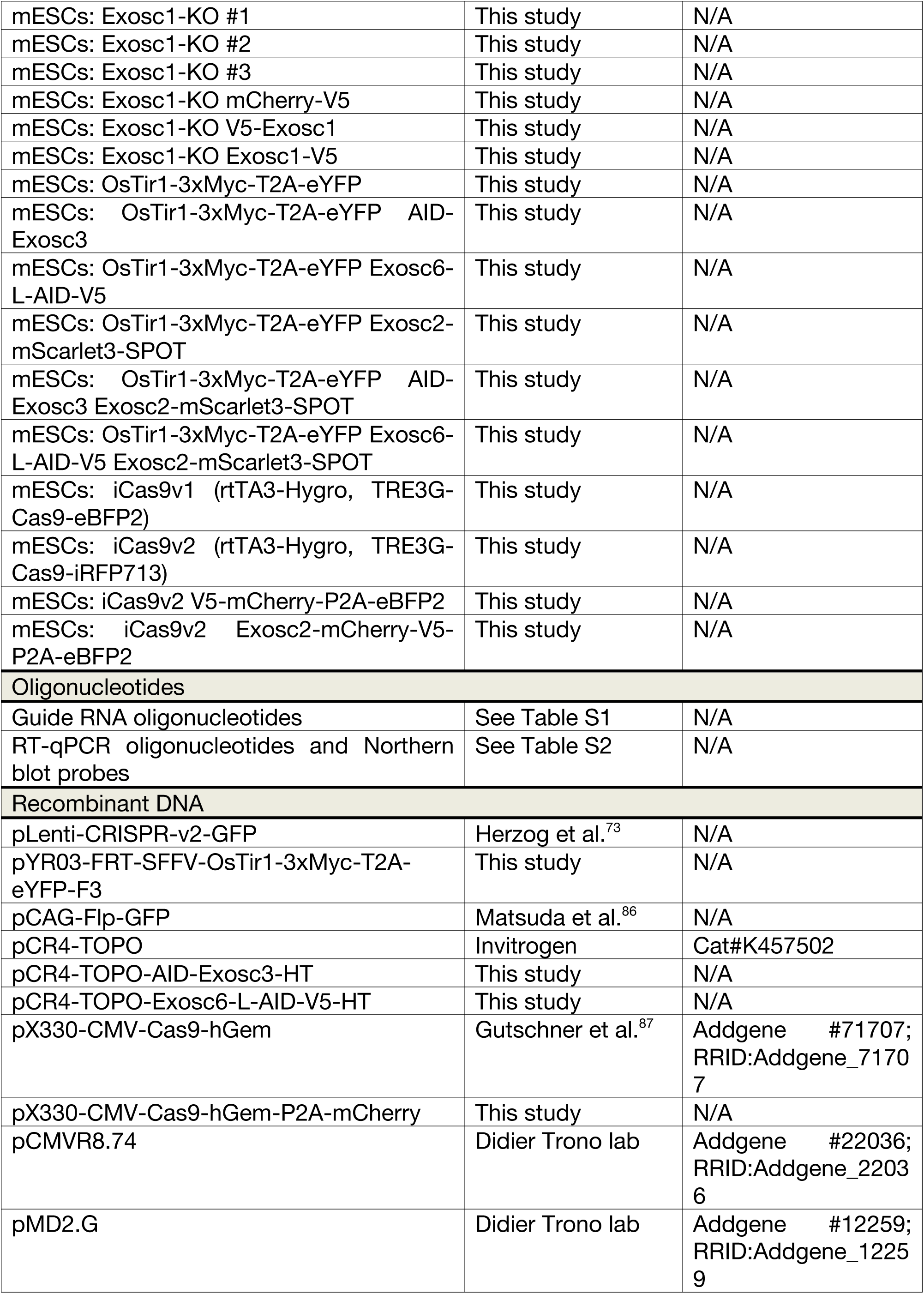
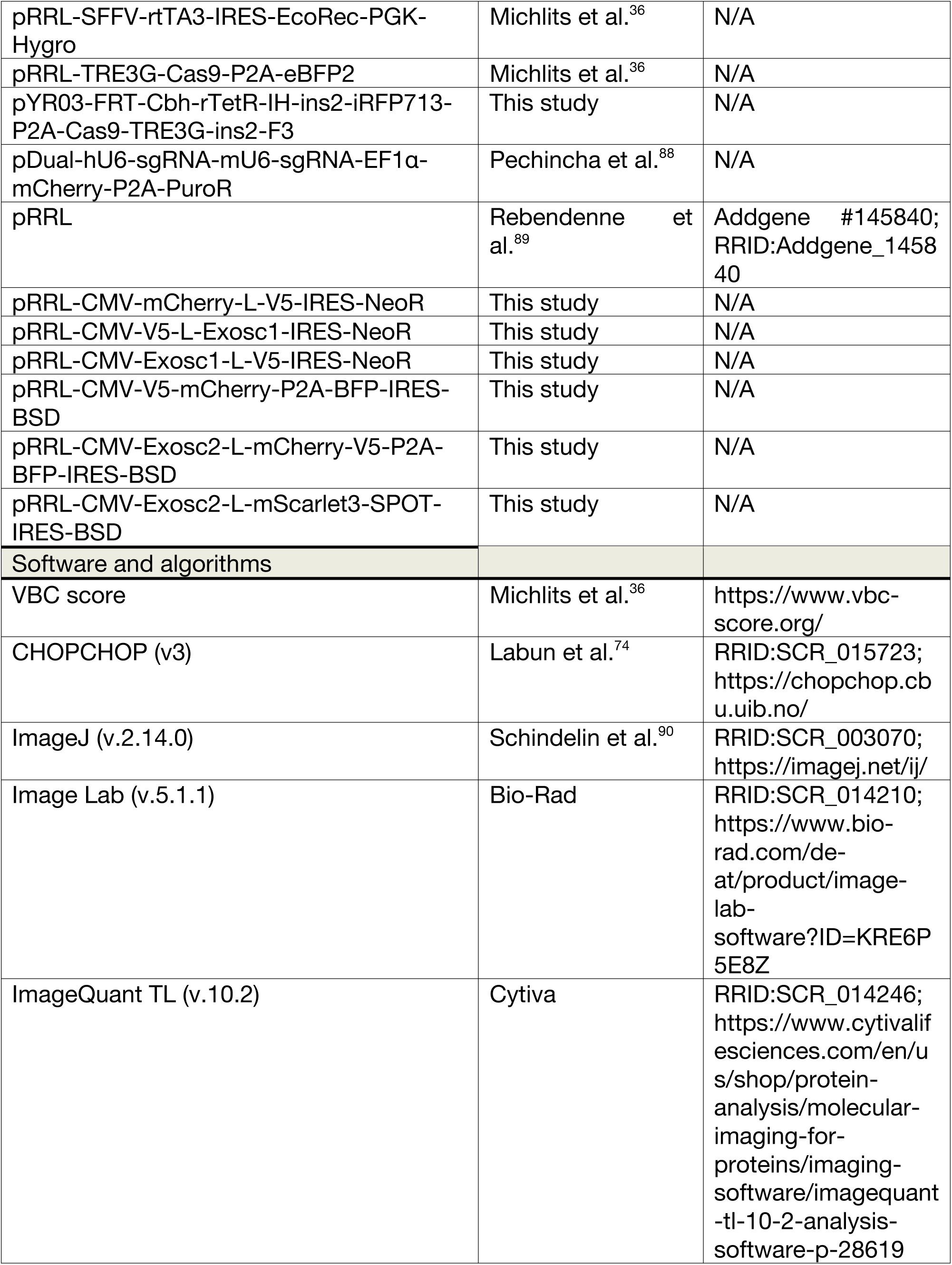
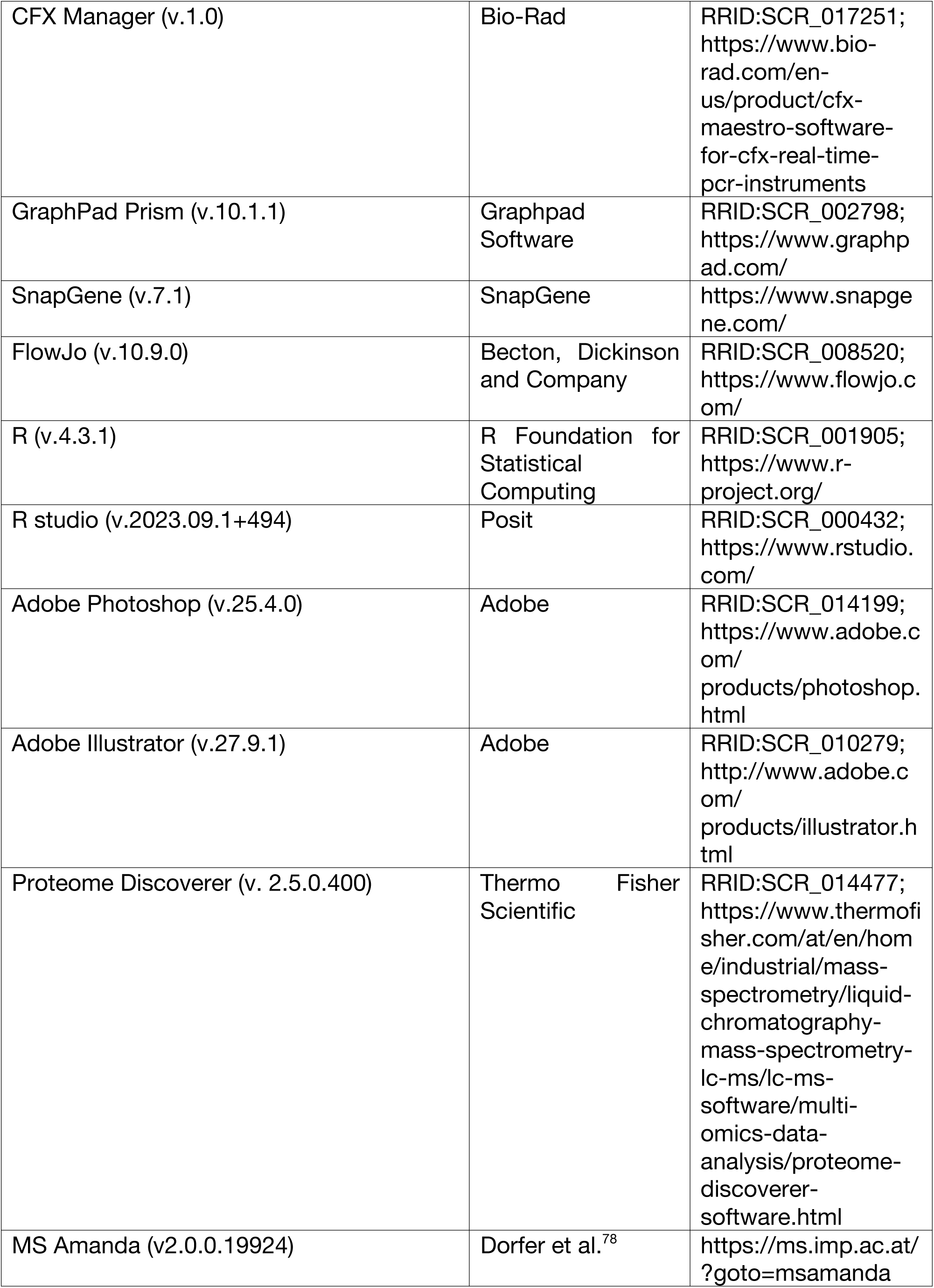
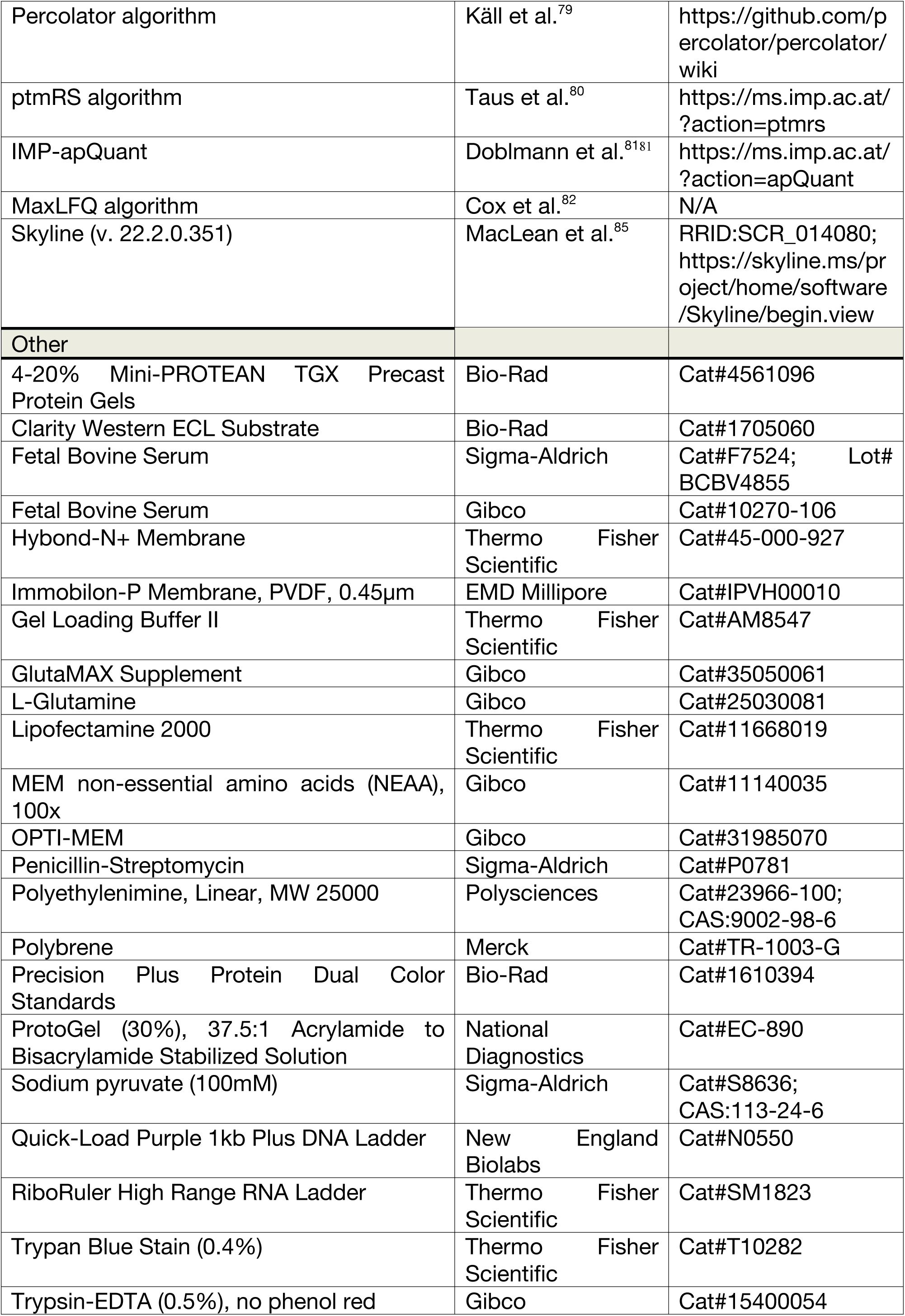

## Supplementary figure titles and legends

**Figure S1.**
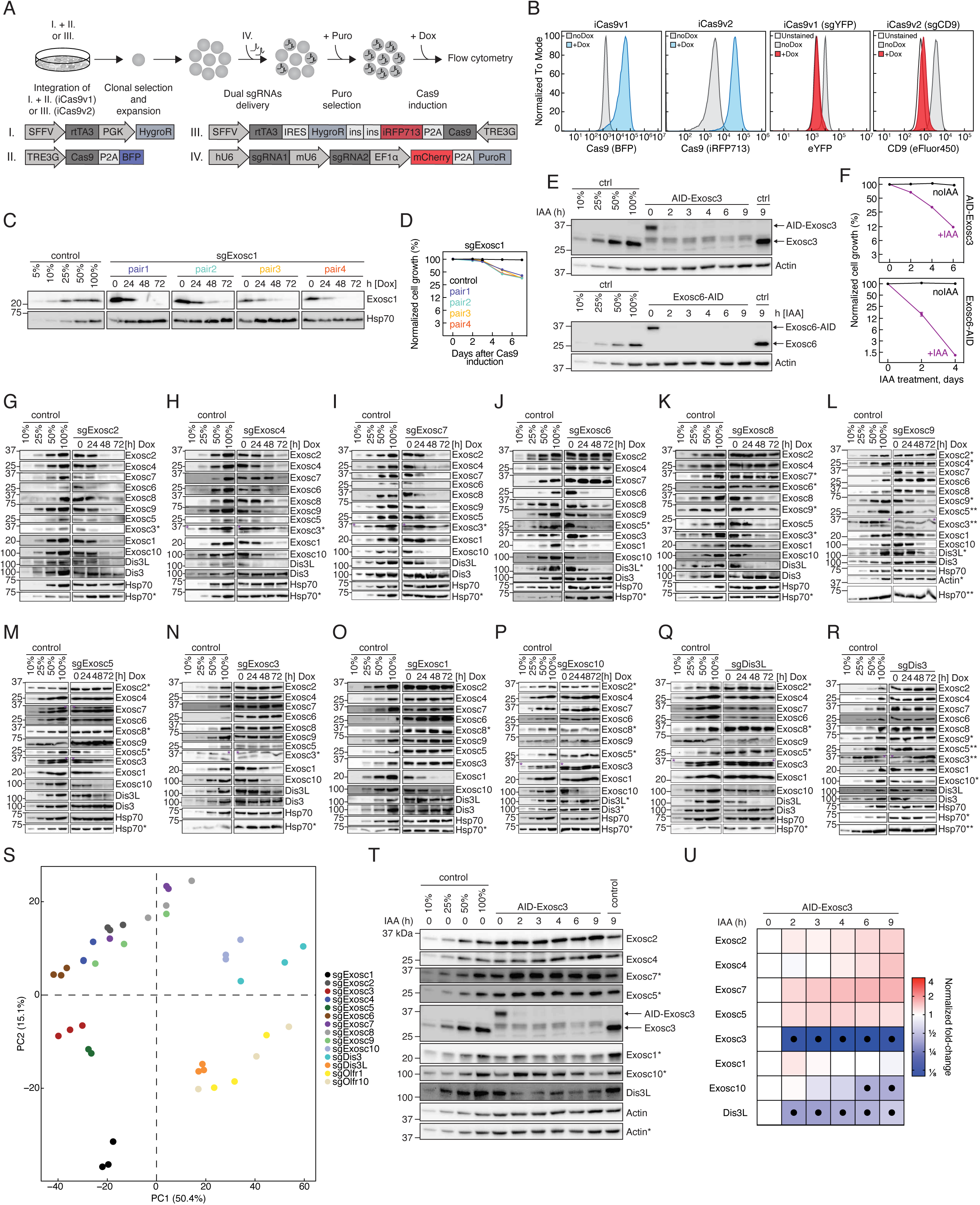
Establishment, validation, and benchmarking of a dual-guide iCas9 system to interrogate proteostatic dependencies among RNA exosome components, related to Figures 1 and 2. **(A)** Schematic of the generation of clonal iCas9 cell lines. Roman numbers indicate vectors used for the engineering of first (iCas9v1) and second (iCas9v2) versions of iCas9. For iCas9v1, expression cassettes for rtTA3 and Cas9 were packaged in lentiviruses and sequentially integrated in the genome. For iCas9v2 “all-in-one” cassette was site-specifically integrated in the expression stable locus on Chr15 (see Materials & Methods). **(B)** (Left): Flow cytometry evaluation of inducible Cas9 expression in iCas9v1 and iCas9v2 in the presence or absence of Dox. (Right): Evaluation of the editing efficiency of endogenously expressed YFP (iCas9v1) or surface marker CD9 (iCas9v2) in iCas9 cells transduced with the respective sgRNAs. For CD9 analysis cells were immunostained before flow cytometry measurements. **(C)** iCas9 cells expressing different pairs of dual sgRNAs against Exosc1 (pair1-4) were treated with Dox for 0h, 24h, 48h and 72h. Total proteins were analyzed with Western blot against Exosc1 and Hsp70. Dilution series (5%, 10%, 25%, 50%, 100%) were used to estimate the sensitivity of the antibody and Hsp70 served as a loading control. **(D)** Competitive proliferation assays of iCas9 cells harboring dual sgRNAs against Exosc1 (pair1-4) and Olfr10 (ctrl). Percentage of sgRNA+ cells was monitored with a flow cytometry. Values were normalized to Day0 and are represented as mean ± sd (n = 2 technical replicates). **(E)** Evaluation of the depletion efficiency of AID-tagged proteins upon IAA administration in AID-Exosc3 (top) and Exosc6-AID (bottom) cells. Tir1 (control) is a parental cell line that served as a negative control. Total proteins were analyzed with Western blot using the indicated antibody. Dilution series (10%, 25%, 50%, 100%) were used to estimate the sensitivity of the antibody and Actin represents a loading control. **(F)** Competitive proliferation assay of AID-Exosc3 (top) and Exosc6-AID (bottom) in the presence and absence of IAA. Percentage of AID-positive cells was monitored with flow cytometry. Values were normalized to Day0 and are represented as mean ± sd (n = 2 biological replicates). **(G-R)** iCas9 cells harboring dual sgRNAs against Exosc2 **(G)**, Exosc4 **(H)**, Exosc7 **(I)**, Exosc6 **(J)**, Exosc8 **(K)**, Exosc9 **(L)**, Exosc5 **(M)**, Exosc3 **(N)**, Exosc1 **(O)**, Exosc10 **(P)**, Dis3L **(Q)** and Dis3 **(R)** were treated with Dox for 0h, 24h, 48h and 72h. Total proteins were analyzed with Western blot for the indicated RNA exosome components. Dilution series (10%, 25%, 50%, 100%) were used to estimate the sensitivity of the antibody and Hsp70 served as a loading control. Black asterisk (*) refers to the corresponding gel and purple asterisk (**\***) indicates a non-specific band. **(S)** Principal component analysis of the mass-spectrometry data from the cells with inducible knockout (72h on Dox) of Olfr1, Olfr10, Exosc1-Exosc10, Dis3 and Dis3L. Olfr1 and Olfr10 served as negative controls. **(T)** Parental Tir1 (ctrl) and AID-Exosc3 cells were treated with IAA for the indicated period of time. Total proteins were analyzed with Western blot using the indicated antibody. Dilution series (10%, 25%, 50%, 100%) were used to estimate the sensitivity of the antibody and Actin served as a loading control. Asterisk (*) refers to the corresponding gel. **(U)** Heatmap of Western blot quantifications for AID-Exosc3 from **(S)**. Protein abundance was normalized to Actin and mean values are shown relative to untreated AID-Exosc3 sample (n = 3 biological replicates). Statistical significance of the difference between a given sample and the untreated AID-Exosc3 control was determined with a two-way ANOVA test and Holm-Šídák multiple comparison correction. Black dots indicate instances with p_adj_ < 0.05.

**Figure S2.**
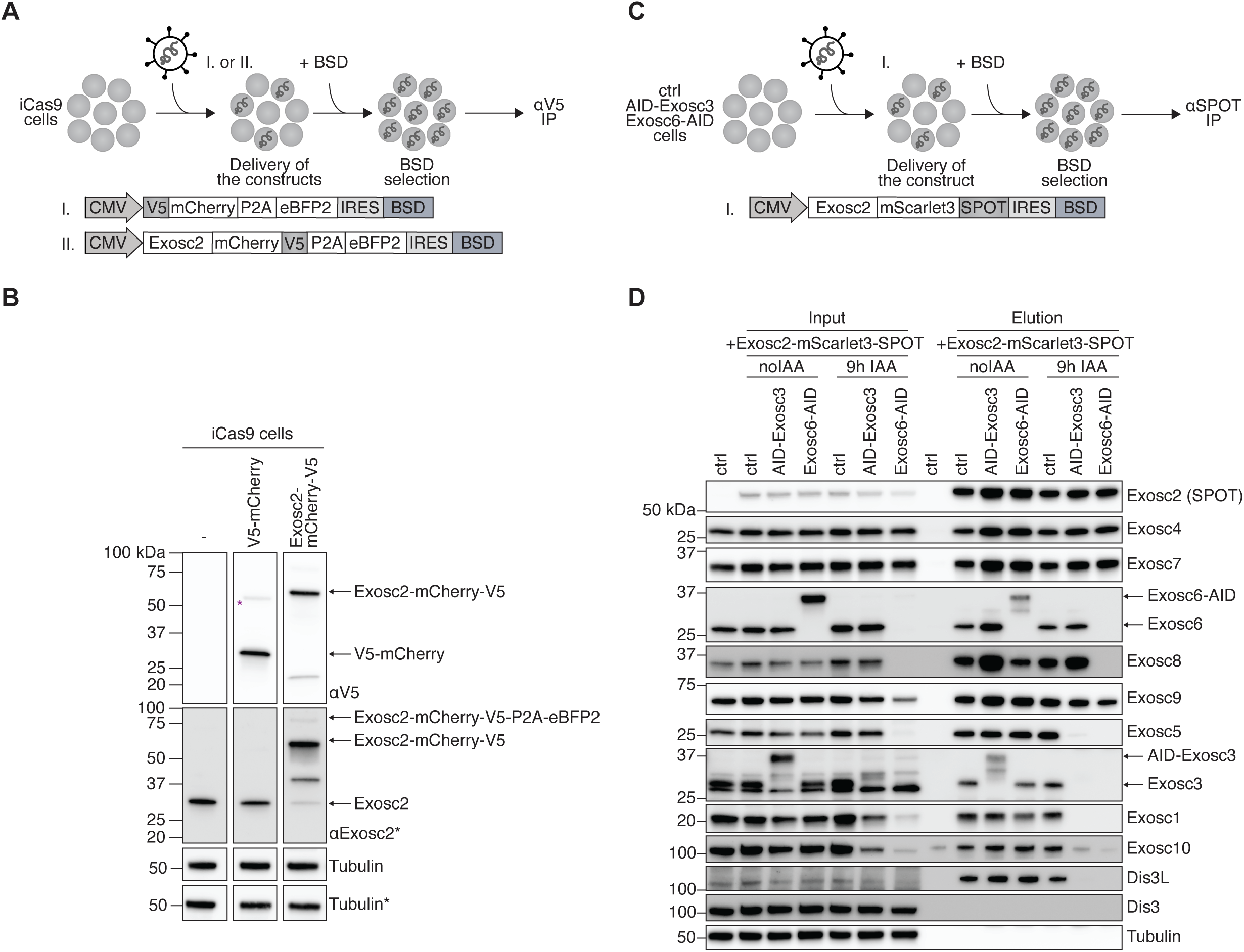
Interrogating RNA exosome assembly intermediates, related to Figure 3. **(A)** Control V5-mCherry-P2A-eBFP2 or Exosc2-mCherry-V5-P2A-eBFP2 cassettes were lentivirally delivered to iCas9 cells followed by BlasticidinS (BSD) selection. **(B)** Western blot analysis of the cell lines from **(A)** with αV5 and αExosc2 antibody. Tubulin served as a loading control. Black asterisk (*) refers to the corresponding gel. Purple asterisk (*) indicates the signal from V5-mCherry-P2A-eBFP2. **(C)** Generation of ectopic expression of Exosc2-mScarlet3-SPOT in the parental Tir1 (ctrl), AID-Exosc3 and Exosc6-AID cell lines. The cassette was virally delivered, and cells were selected with BlasticidinS (BSD). **(D)** αSPOT-IP of cells from **(C)**. Input and elution samples were analyzed with Western blot using the indicated antibody. Tubulin served as a loading control.

**Figure S3.**
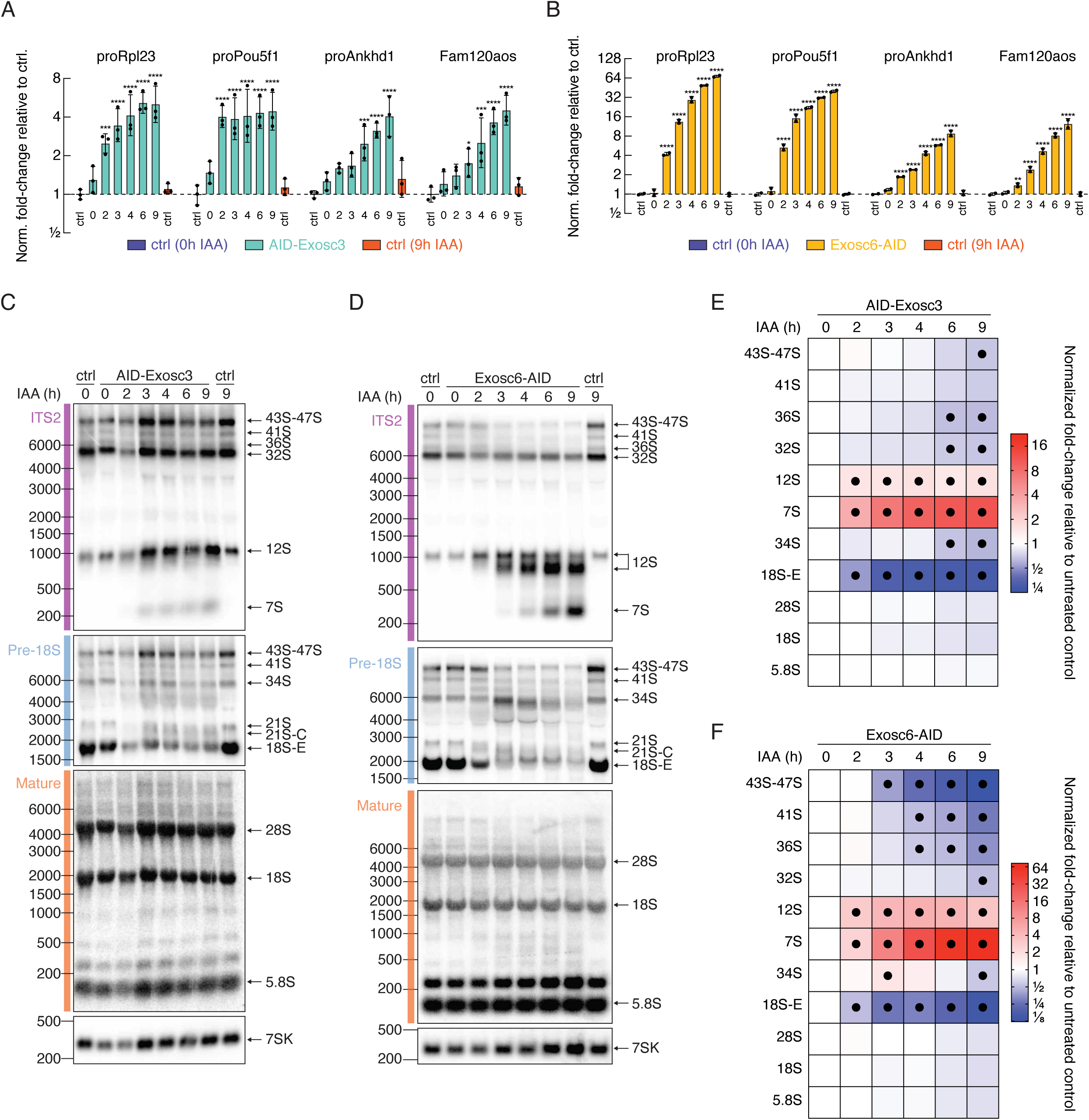
Derepression of PROMPTs and accumulation of rRNA processing defects upon depletion of Exosc3 and Exosc6 by AID, related to Figure 4. **(A-B)** Parental Tir1 (ctrl), AID-Exosc3 **(A)** and Exosc6-AID **(B)** cells were treated with IAA for the indicated time period. Total RNA was isolated and PROMPTs abundance was measured with RT-qPCR. Values were normalized to Gapdh mRNA and mean ± sd is shown relative to ctrl 0h IAA (n = 3 for AID-Exosc3 and n=2 for AID-Exosc6 biological replicates). Statistical significance of the difference between a given sample and “0h IAA ctrl” was determined with a two-way ANOVA test and Holm-Šídák multiple comparison correction (* p_adj_ < 0.05, ** p_adj_ < 0.0021, *** p_adj_ < 0.0002, **** p_adj_ < 0.0001, non-significant p_adj_ > 0.05 is not marked). **(C-D)** Parental Tir1 (ctrl), AID-Exosc3 **(C)** and Exosc6-AID **(D)** cells were treated with IAA for the indicated time period. Total RNA was resolved on the agarose gel followed by northern blot analysis. **(E-F)** Heatmap of northern blot quantifications for AID-Exosc3 **(E)** and Exosc6-AID **(F)**. Abundance of rRNA processing intermediates was normalized to 7SK RNA and mean is shown relative to untreated AID-Exosc3 or Exosc6-AID samples (n = 3 for AID-Exosc3 and n=2 for Exosc6-AID biological replicates). Statistical significance of the difference between a given sample and untreated AID-Exosc3 or Exosc6-AID controls was determined with a two-way ANOVA test and Holm-Šídák multiple comparison correction. Black dots indicate instances with p_adj_ < 0.05.

**Figure S4.**
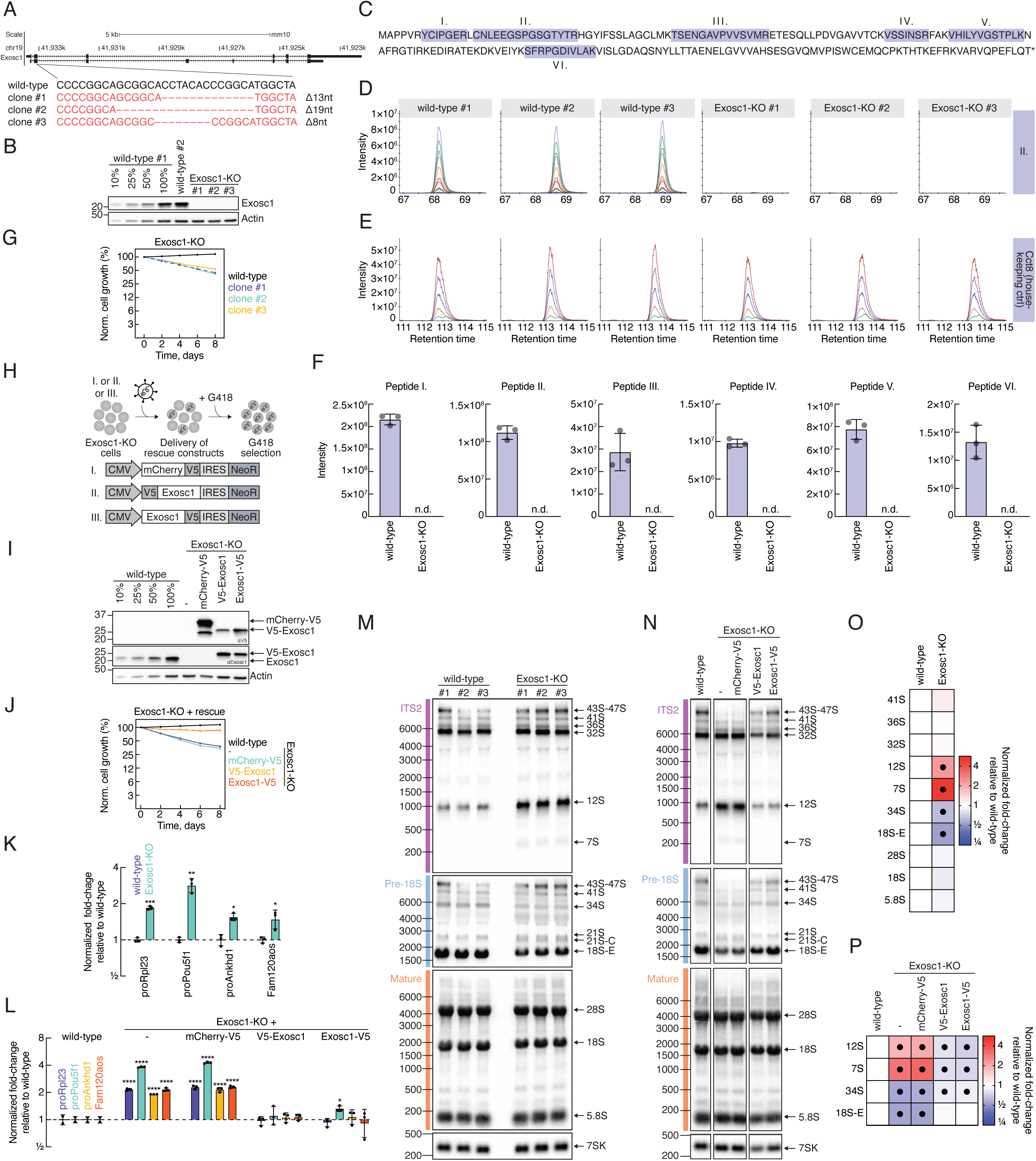
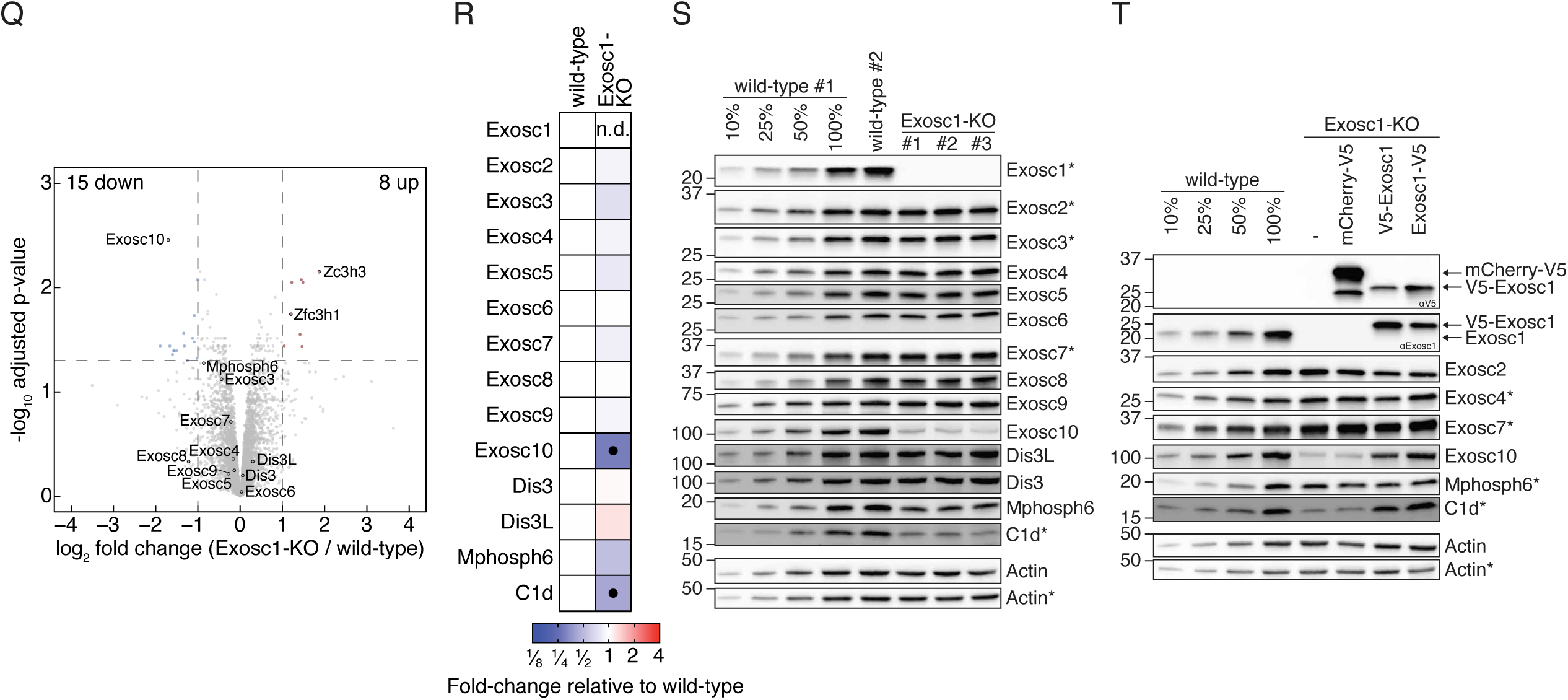
Establishment, validation and characterization of mESC lines depleted of Exosc1 by CRISPR/Cas9 genome engineering, related to Figure 4. **(A)** Schematic of Exosc1 genomic loci. Frameshift mutations were introduced in the second coding exon with CRISPR/Cas9 system. Genotypes of three Exosc1-KO clones used in this study are indicated. **(B)** Western blot analysis of wild-type and clonal Exosc1-KO cell lines. Dilution series (10%, 25%, 50%, 100%) were used to estimate the sensitivity of the antibody and Actin represents a loading control. **(C)** Protein sequence of Exosc1. Six peptides that were measured with parallel reaction monitoring (PRM) mass-spectrometry are highlighted with blue rectangles and labelled with roman numbers (I-VI). **(D)** Example PRM analysis of wild-type and Exosc1-KO cells for the peptide II. PRM intensity signal (y-axis) is plotted against the retention time (x-axis). Multiple curves correspond to MS/MS=MS2 fragments. **(E)** Example PRM analysis of one of the peptides of Cct8 protein that served as a “loading control”. Data is represented as in **(D)**. See Materials & Methods for the information about the reference proteins used in PRM experiments. **(F)** PRM signal intensity of Exosc1 peptides in wild-type and Exosc1-KO cells. No signal was detected in Exosc1-KO samples and is depicted as n.d. (not determined). **(G)** Competitive proliferation assay of wild-type and Exosc1-KO cells. Percentage of wild-type or Exosc1-KO cells was monitored with flow cytometry. Values were normalized to Day0 and are shown as mean ± sd (n = 2 technical replicates). **(H)** Schematic of Exosc1 rescue experiments. Rescue constructs harboring mCherry-V5 or V5-Exosc1 or Exosc1-V5 were virally delivered to Exosc1-KO cells followed by G418 selection. **(I)** Western blot analysis of the cells with rescue constructs. Membranes were probed with αV5 and αExosc1 antibody to confirm transgene expression. Dilution series (10%, 25%, 50%, 100%) were used to estimate the sensitivity of the antibody and Actin served as a loading control. **(J)** Competitive proliferation assay, as in **(G)**, for Exosc1 rescue cells. **(K-L)** Total RNA from wild-type, Exosc1-KO **(K)** and rescue **(L)** experiments was isolated and PROMPTs abundance was measured with RT-qPCR. Values were normalized to Gapdh mRNA and mean ± sd is shown relative to wild-type (n = 3 biological replicates). Statistical significance of the difference between a given sample and wild-type was determined with unpaired t-test (for **K**) or two-way ANOVA (for **L**) and Holm-Šídák multiple comparison correction. (n = 3 biological replicates, (* p_adj_ < 0.05, ** p_adj_ < 0.0021, *** p_adj_ < 0.0002, **** p_adj_ < 0.0001, non-significant p_adj_ > 0.05 is not marked). **(M-N)** Total RNA from wild-type, Exosc1-KO **(M)** and rescue **(N)** experiments was resolved on the agarose gel followed by northern blot analysis. **(O-P)** Heatmap of northern blot quantifications for wild-type, Exosc1-KO **(O)** and rescue **(P)** experiments. Values were normalized to 7SK RNA and mean is presented relative to wild-type (n = 3 biological replicates). Statistical significance of the difference between a given sample and wild-type was determined with unpaired t-test (for **O**) or two-way ANOVA (for **P**) and Holm-Šídák multiple comparison correction (n = 3 biological replicates. Black dots indicate instances with p_adj_ < 0.05. (R) Volcano plots displaying whole proteome changes for Exosc1-KO. The log_2_ fold change (x axis) is plotted against the -log10 of Benjamini–Hochberg adjusted limma p-value (y axis; p_adj_). Dashed lines indicate the thresholds of p_adj_ < 0.05 and log_2_FC < -1 or > 1 (n = 3 biological replicates). (S) Relative protein abundance of RNA exosome subunits in Exosc1-KO determined by shotgun and PRM mass-spectrometry. Values are shown relative to wild-type sample. Statistical significance of the difference between Exosc1-KO and wild-type was determined with an unpaired t-test and Holm-Šídák multiple comparison correction (n = 3 biological replicates). Black dots indicate instances where p_adj_ < 0.05. No signal for Exosc1 peptides was measured in Exosc1-KO samples and is depicted as n.d. (not determined). See also panel **(F)**. **(S-T)** Western blot analysis of wild-type, Exosc1-KO **(S)** and rescue cells **(T)** with the indicated antibody. Dilution series (10%, 25%, 50%, 100%) were used to estimate the sensitivity of the antibody and Actin represents a loading control. Asterisk (*) refers to the corresponding gel.

